# Spatiotemporal optical control of Gαq-PLCβ interactions

**DOI:** 10.1101/2023.08.10.552801

**Authors:** Sithurandi Ubeysinghe, Dinesh Kankanamge, Waruna Thotamune, Dhanushan Wijayaratna, Thomas M. Mohan, Ajith Karunarathne

## Abstract

Cells experience time-varying and spatially heterogeneous chemokine signals in vivo, activating cell surface proteins, including G protein-coupled receptors (GPCRs). The Gαq pathway activation by GPCRs is a major signaling axis with a broad physiological and pathological significance. Compared to other Gα members, GαqGTP activates many crucial effectors, including PLCβ (Phospholipase Cβ) and Rho GEFs (Rho guanine nucleotide exchange factors). PLCβ regulates many key processes, such as hematopoiesis, synaptogenesis, and cell cycle, and is therefore implicated in terminal - debilitating diseases, including cancer, epilepsy, Huntington’s Disease, and Alzheimer’s Disease. However, due to a lack of genetic and pharmacological tools, examining how the dynamic regulation of PLCβ signaling controls cellular physiology has been difficult. Since activated PLCβ induces several abrupt cellular changes, including cell morphology, examining how the other pathways downstream of Gq-GPCRs contribute to the overall signaling has also been difficult. Here we show the engineering, validation, and application of a highly selective and efficient optogenetic inhibitor (Opto-dHTH) to completely disrupt GαqGTP-PLCβ interactions reversibly in user-defined cellular-subcellular regions on optical command. Using this newly gained PLCβ signaling control, our data indicate that the molecular competition between RhoGEFs and PLCβ for GαqGTP determines the potency of Gq-GPCR-governed directional cell migration.

## 1. Introduction

G protein-coupled receptors (GPCRs) are a large family of proteins encoded by ∼5% of human genes^1, 2^. In humans, approximately a thousand GPCRs are widely expressed in various tissues, controlling various physiological processes^3^. Therefore, many diseases involve dysregulated GPCR signaling, making them major drug targets^3^. The enormous diversity of GPCRs allows them to sense various ligands such as odorants, neurotransmitters, and hormones^4, 5^. GPCRs activate a diverse group of heterotrimeric G proteins comprising Gα and Gβγ subunits. There are 16Gα, 5Gβ, and 12Gγ types in humans,^6^ in which the 16Gα subunits are classified into four major families with distinct signaling; Gαs, Gαi/o, Gαq/11, and Gα12/13^7^.

Activated GPCRs serve as Guanine nucleotide Exchange Factors (GEFs) for Gα subunits and facilitate GDP to GTP exchange on the interacting GαGDPβγ heterotrimer, resulting in GαGTP and free Gβγ^8^. Upon activation, GαqGTP activates phospholipase C β (PLCβ) isoforms (PLCβ1,2,3 and 4) to different extents^9, 10^, inducing the hydrolysis of the inner membrane lipid phosphatidylinositol 4,5-bisphosphate (PIP2) into diacylglycerol (DAG) and inositol 1,4,5-triphosphate (IP3)^11^. GαqGTP activates PLCβ1 and 3 to a greater extent than PLCβ2 and 4^10^. The resultant IP3 induces mobilization of stored calcium and regulates cell proliferation and other cellular functions that require calcium. DAG activates protein kinase C (PKC) and promotes immune cell activation and regulation^11–13^. Dysregulated IP3 signaling is involved in heart and brain diseases such as neurodegenerative diseases and ischemia,^14^ while DAG is implicated in visceral obesity, insulin resistance, and atherosclerosis. GαqGTP also activates Rho guanine nucleotide exchange factors (GEFs), which catalyze the GDP to GTP exchange on RhoA. Subsequently, RhoA activates Rho kinase 1/2 (ROCK 1/2), and mDia, catalyzing myosin light change phosphorylation and subsequent actomyosin contractility^15^. RhoA signaling is involved in several pathological conditions, including neurodegenerative diseases such as Alzheimer’s, Parkinson’s, amyotrophic lateral sclerosis, and Huntington’s disease^16^.

The structural data shows that PLCβ3 interacts with GαqGTP via three primary contact points^10^. The first involves Gαq switch residues, and residues in the loop between PLCβ 3/4 EF-hand domains^10^. Switch regions I and II of Gαq and the C2 domain and the loop connecting it to the catalytic TIM barrel of PLCβ3 form the second contact point^10^. The third contact site in Gαq comprises the interactions between the switch region II and the neighboring α3 domain of GαqGTP and the conserved helix-turn-helix (HTH) region of the PLCβ3^10, 11, 17^. Among the three, the conserved HTH region of PLCβ3 serves as the major surface for PLCβ3-Gαq interactions^10, 11^. TAMRA-labeled peptides representing the complete HTH domain of PLCβ3 from *Tyr*847 to *Glu*882 have exhibited interactions with purified and aluminum fluoride (AlF^−^)-activated Gαq^18^. AlF^−^ interacting GαGDP is expected to have a structure analogous to GαGTP^19^, which led to the conclusion that HTH peptide forms robust interactions with GαqGTP, preventing the engagement and activation of GαqGTP downstream effectors. In particular, HTH residues; *Tyr*855, *Leu*859, *Asn*861, *Pro*862, *Ile*863, and *Asp*870 have shown strong interactions with the groove formed between switch II and helix α3 in active Gαq^19^. Further, the ability of the HTH domain to inhibit Gαq signaling has been investigated *in vivo* ^19^. Injection of purified HTH peptides into mice prefrontal cortex neurons has shown a small yet significant inhibition of Gαq-triggered depolarization^19^. Interestingly, these peptides showed no interactions with GαqGDP or other GαGTPs^19^.

PLCβ signaling involves many pathological conditions, including brain disorders, hematologic diseases, impaired T-cell migration, and cancers^20^. However, the lack of pharmacological inhibitors perturbs understanding of how dynamically PLCβ is implicated in pathology. Additionally, since GαqGTP is a major disease-driving signaling axis, segregation of PLCβ signaling from other Gαq-effector signaling can help isolate beneficial and disease-driving components of the Gαq pathway. To achieve these, we developed a series of optogenetic inhibitors for PLCβ using HTH substructure of PLCβ3 to exclusively and reversibly disrupt GαqGTP - PLCβ interaction with high efficacy. We recruited *Avena sativa* Light-Oxygen-Voltage sensing domain (AsLOV2)-based iLID (improved light-induced dimer) and the small protein, SspB dimerization upon blue light^21–24 25^ to present optimized peptides to GαqGTP on optical command to disrupt GαqGTP-PLCβ interactions. Our optogenetic inhibitor exclusively and reversibly disrupts GαqGTP-mediated PLCβ in user-defined single cells and subcellular regions. The sample findings show how PLCβ influences RhoA signaling and regulates the Gαq-pathway-governed cell migration.

## 2. Results and Discussion

### 2.1 Optogenetic inhibition feasibility of GαqGTP-PLCβ signaling

The switch region II and the neighboring α3 domain of GαqGTP interact with the conserved helix-turn-helix (HTH) region of the PLCβ3^26^ (Fig. 1A). We first examined whether the cytosolic HTH peptide from *Tyr*847 to *Glu*882 could inhibit Gαq-coupled Gastrin Releasing Peptide Receptor (GRPR) induced PIP2 hydrolysis in HeLa cells. We used the Venus-tagged PH domain of PLCδ1 to measure PIP2 hydrolysis, which translocates from the plasma membrane to the cytosol upon PIP2 hydrolysis^27^. We recently showed that Gαq-pathway induced PIP2 hydrolysis is a transient and self-attenuating process governed by the dissociation of Gβγ from the initially formed GαqGTP–PLCβ–Gβγ sandwich complex^11^. This self-attenuation is indicated by the ∼50% PIP2 hydrolysis reduction (Fig. 1B, blue arrow). Here, HeLa cells expressing either cytosolic mRFP (control) or mRFP-HTH showed typical and similar PIP2 hydrolysis-attenuation responses upon GRPR activation with 1 μM of bombesin (Fig. 1B). This indicated that the cytosolic HTH does not perturb GαqGTP-PLCβ signaling, and the mean maximum extents of PIP2 hydrolysis between the two were similar (one-way ANOVA*: F*_1, 38_ =0.03339, *p* =0.85598; Fig. S1A, Supplemental Table S2, A, and B). We next examined whether a plasma membrane-targeted HTH could inhibit Gαq-induced PIP2 hydrolysis. Compared to the control cells expressing membrane-targeted mRFP (Lyn-mRFP), the cells expressing an N terminally palmitoylated HTH (Lyn-mRFP-HTH) did not show a detectable PIP2 hydrolysis response upon GRPR activation (Fig. 1C). This indicated that when the HTH domain is spatially proximal, it can effectively disrupt PLCβ activation. We then examined whether membrane-targeted HTH also inhibits GαiGTP or GαsGTP-governed signaling. HeLa cells expressing Blue opsin-mTurquoise (BO-MTQ, a Gi/o-GPCR), cAMPr (cAMP sensor), and Lyn-mRFP-HTH (test), or Lyn-mRFP (control) were imaged using 488 nm excitation/515 nm emission to capture cAMPr fluorescence at 1-sec intervals. At 30 sec, we activated adenylyl cyclase by exposing cells to 10 μM forskolin in 20 μM IBMX (3-isobutyl-1-methylxanthine), which increased cAMPr fluorescence (Fig. S1B)^28^, and at 200 s, 5 μM 11-cis-retinal (final concentration in cells) was added. Since we imaged cells using 488 nm, retinal addition activated blue opsin, and both cell types (Lyn-mRFP-HTH and Lyn-mRFP) exhibited similar reductions in cAMPr fluorescence (Fig. S1B). Control cells without 11-cis-retinal exposure showed a sustained cAMPr fluorescence after adenylyl cyclase activation (Fig. S1B-plot-dash lines). This data indicated that plasma membrane-bound HTH doesn’t perturb GαiGTP-mediated adenylyl cyclase inhibition. Similarly, GαsGTP-adenylyl cyclase-induced cAMP generation was examined in HeLa cells expressing β1AR-CFP (a Gs-GPCR), cAMPr, and Lyn-mRFP-HTH (test) or Lyn-mRFP (control) (Fig. S1C). Both control and test cells showed similar cAMPr fluorescence increases upon addition of 10 μM isoproterenol, indicating that HTH doesn’t disrupt GαsGTP-adenylyl cyclase interactions.

**Figure. 1:**
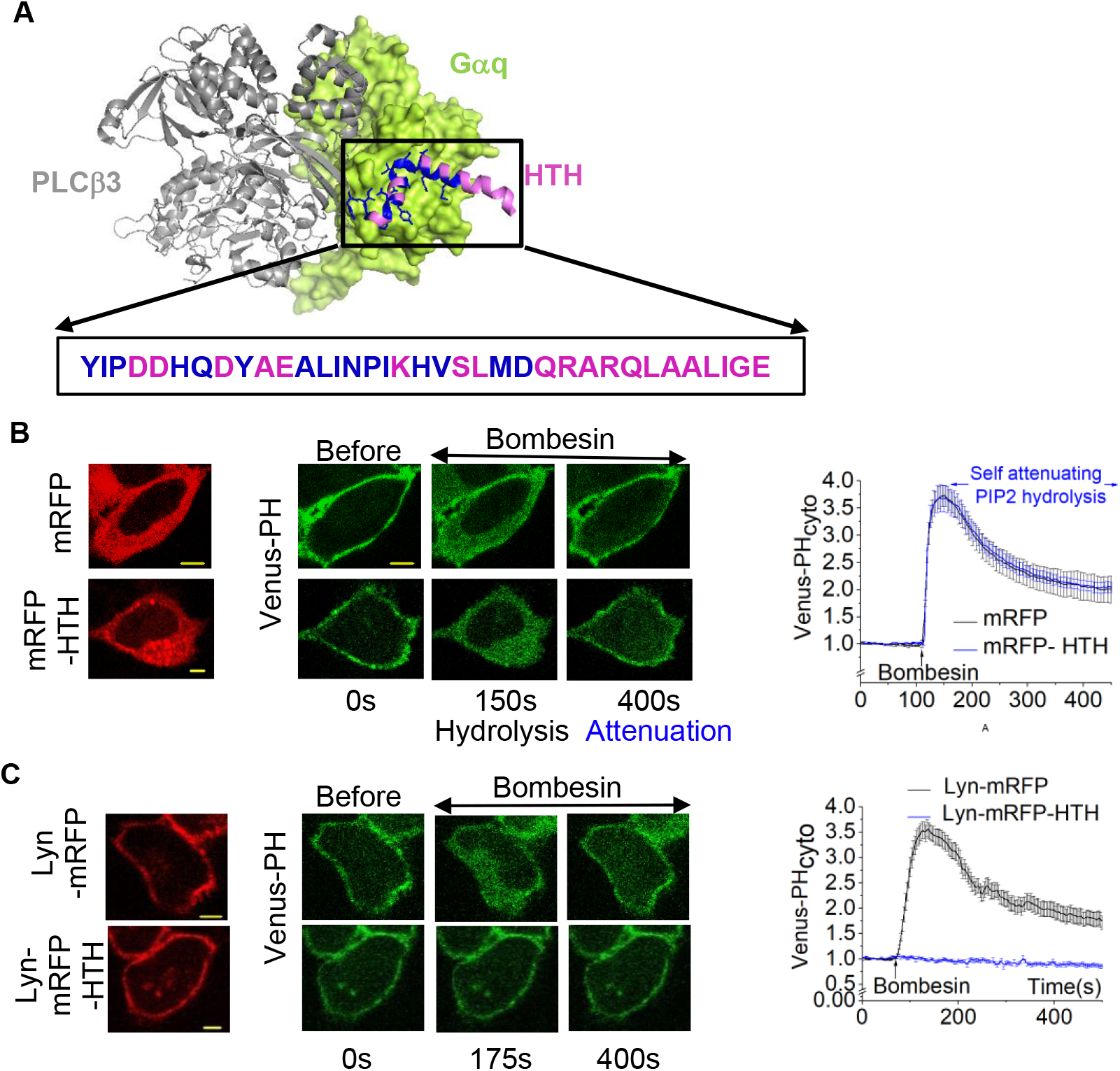
PLCβ3 HTH domain blocks the PLCβ binding site on GαqGTP and inhibits the downstream PLCβ signaling. **(A)** Diagram depicting the interactions between PLCβ3 and Gαq. HTH domain (pink) of PLCβ3 (grey) interacts with the Gαq (light green). The HTH residues that form strong interactions with GαqGTP are shown in blue color. **(B)** HeLa cells expressing GRPR, Venus-PH, and cytosolic mRFP-HTH or mRFP exhibited efficient PIP2 hydrolysis upon adding 1 μM bombesin (n =18 for each condition). The corresponding plot shows the PIP2 sensor dynamics in the cytosol of the cells. **(C**) In contrast to the HeLa cells expressing GRPR, Venus-PH, and Lyn-mRFP (control), the cells expressing GRPR, Venus-PH, and Lyn-mRFP-HTH didn’t show significant PIP2 hydrolysis upon activation with 1 μM bombesin. The corresponding plot shows the PIP2 sensor dynamics in the cytosol of the cells (n =16 for each condition). The error bars represent SEM (standard error of the mean). The scale bar = 5 µm. GRPR: Gastrin Releasing Peptide receptor; mRFP: Monomeric Red Fluorescent protein; PM: Plasma membrane; PIP2: Phosphatidylinositol 4,5-bisphosphate; Cyto: cytosolic fluorescence; HTH: Helix-Turn-Helix; PH: Pleckstrin Homology.

To understand whether HTH broadly inhibits GαqGTP signaling or selectively prevents PLCβ activation, we examined whether HTH perturbs GαqGTP-RhoGEFs interactions. We expressed GRPR, mTurquoise-PH, and Venus-rGBD (a RhoA sensor) that translocates from the cytosol to the plasma membrane upon RhoA activation,^29^ together with either Lyn-mRFP-HTH (test) or Lyn-mRFP (control). Upon GRPR activation in Lyn-mRFP-HTH cells, although mTurquoise-PH remained on the plasma membrane indicating PIP2 hydrolysis inhibition by HTH (Fig. S1D-left panel), a pronounced Venus-rGBD translocation from the cytosol to the plasma membrane was observed due to endogenous RhoA activation (Fig. S1D-left panel). This indicates that although membrane-targeted HTH completely inhibits GαqGTP-PLCβ interactions, GαqGTP can still activate RhoGEFs in HeLa cells. Nevertheless, not only cells expressing Lyn-mRFP showed PIP2 hydrolysis, their Venus-rGBD recruitment to the plasma membrane was significantly lower than that of Lyn-mRFP-HTH cells (one-way ANOVA*: F*_3, 12_ =377.51099, *p* =<0.0001; Fig. S1D-plot, Supplemental Table S3, A, and B). The reduced RhoA activation here is likely due to the compromised ability of RhoGEFs to interact with PLCβ-bound GαqGTP over the HTH-bound in Lyn-mRFP-HTH cells (Fig. S1D- right panel). Although we have not determined whether HTH only permits specific RhoGEF activities, we propose that it will only perturb activities of RhoGEFs with HTH domain, such as p63RhoGEF. This data can be reasoned since our RNA seq data and homology modeling also show that HeLa cells primarily express RhoGEFs lacking the HTH domain, including LARG and TRIO (Fig. S1E and table S4).

Since GαqGTP activates several PLCβ isoforms and the position weight matrix shows that the amino acid sequences of all PLCβ isoforms possess significantly similar HTH domains (Fig. S2A and B), we built homology models of PLCβ1, 2, and 4 using the available full-length PLCβ3 structure (PDB ID:4GNK)^30^. Considering the predicted significant structural similarity of these HTH domains, we believe that molecular tools based on the HTH domain of PLCβ3 can inhibit the interactions between GαqGTP and all four PLCβ isoforms.

Considering the above distinct, spatial proximity-driven PLCβ signaling inhibitory paradigms of HTH; i. e., no-inhibition by cytosolic and complete inhibition by plasma membrane-targeted HTH (Fig. 1B and C), we next examined the feasibility of optically controlling HTH proximity to the plasma membrane. We used the blue light-dependent dimerization module Cryptochrome 2 (CRY2) and CIBN^31^. CRY2-CIBN module-based optogenetic targeting of proteins has widely been used to control signaling on optical command^31–34^. We generated CRY2-mCherry-HTH and expressed it in HeLa cells with the plasma membrane-targeted CIBN-CAAX, the CRY2 interacting domain^31, 33^. Cytosolic CRY2-mCherry-HTH without blue light exposure did not perturb GRPR-induced PIP2 hydrolysis (Fig. S2C-top panel). Interestingly, although blue light exposure induced robust CRY2-mCherry-HTH plasma membrane recruitment, it also did not inhibit GRPR-induced PIP2 hydrolysis (Fig. S2C-bottom panel). This suggested that HTH in CRY2 does not have sufficient spatial proximity or flexibility, or both, that the plasma membrane-bound Lyn-mRFP-HTH possessed to interact with GαqGTP.

### 2.2 Engineering of iLID-based GαqGTP-PLCβ optogenetic inhibitor

The rationale for using the Improved light-induced dimerization module (iLID) in inhibitor engineering is to enhance the spatial proximity of the inhibitory peptide to GαqGTP upon blue light exposure. Here we utilize iLID’s significantly smaller size than CRY2 (iLID’s 17kDa vs. CRY2’s 58kDa). Dark state PAS domain in iLID contains a non-covalently bound oxidized FMN molecule that absorbs blue light forming a covalent link between FMN and the thiol moiety of the active site *Cys*^22, 35^. This adduct formation drives conformational changes, unwinding the Ct α helix (Jα) upon photoexcitation^21, 22, 24, 36–39^. Blue light-induced unmasking of Jα-integrated short SsrA peptide (-AANDENYF) in iLID allows SsrA to bind its partner, SspB ^25, 40, 41^. We tethered the HTH domain to the Ct of iLID and created Venus-iLID-HTH (Fig.2A-modelled structure and cartoon) and expressed together with the membrane-targeted SspB (Lyn-SspB), GRPR, and the PIP2 sensor-mCherry-PH in HeLa cells. Venus-iLID-HTH showed a cytosolic distribution (Fig. 2B). GRPR activation-induced normalized maximum PIP2 hydrolysis in control cells (without Venus-iLID-HTH expression) and Venus-iLID-HTH expressing cells were similar, indicating that cytosolic HTH has no inhibitory effect (one-way ANOVA: *F_1, 39_* = 1.23122, *p* = 0.27396; Supplemental Table S5A, B, and Fig.S3A). A robust Venus-iLID-HTH recruitment to the plasma membrane was observed upon blue light exposure. Demonstrating a successful optogenetic PLCβ inhibition, GRPR activation only induced minor mCherry fluorescence increase in the cytosol, indicating only a residual PIP2 hydrolysis (Fig.2C-blue box). To comparatively examine the inhibition of GRPR-triggered perpetual interactions between GαqGTP-PLCβ only in selected cells, after GRPR activation, we exposed one of the two cells in the same field of vision to blue light (cell 2) and recruited Venus-iLID-HTH to the plasma membrane (Fig.S3B). The control cell (cell 1) shows the classical PIP2 hydrolysis and its attenuation (Fig. S3B-Plot, black curve). We have recently described that this attenuation is due to the Gβγ translocation away from the plasma membrane^11^. In cell 2, recruitment of Venus-iLID-HTH to the membrane instantaneously disrupted PIP2 hydrolysis as indicated by the 30X faster recovery of PIP2 (PIP2 hydrolysis attenuation rates; Cell 1:6.43×10^−3^ s^−1^ < Cell 2: 2.15 × 10^−2^ s^−1^) (Fig.S3B, blue plot) determined as described previously^11^. Interestingly, once the blue light is terminated on cell 2, a minor yet significant rescue of PIP2 hydrolysis occurred that reached the control cell-level (black arrow on the plot). This also indicated that the HTH-induced inhibition is reversible and only requires blue light termination (Fig. S3B-yellow arrow). The reduced nature of the PIP2 hydrolysis recovery upon blue light termination can be understood by comparatively examining the control cell (cell 1) response, which showed hydrolysis attenuation following the mechanism we described previously ^11^.

**Figure. 2:**
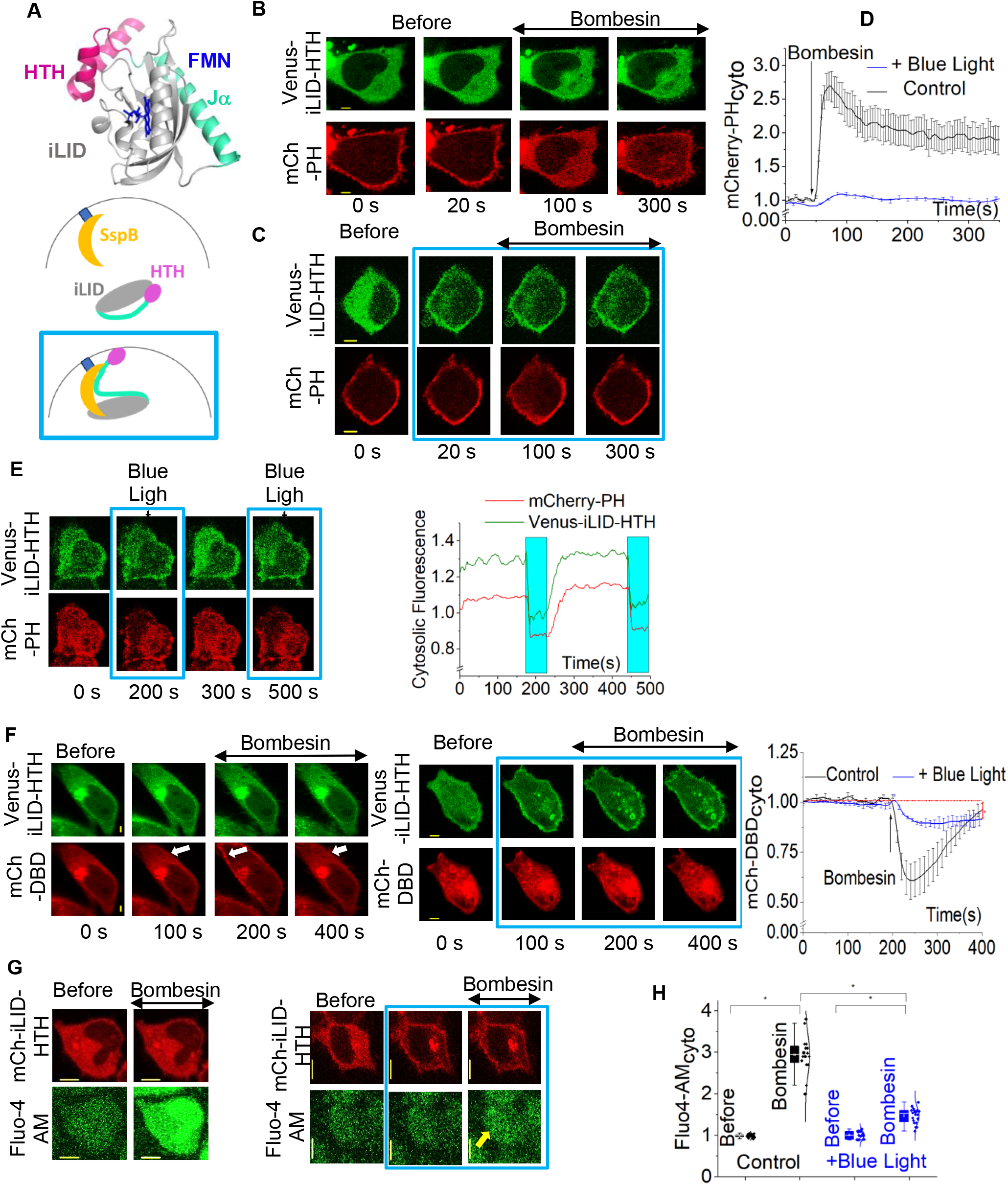
iLID-based optogenetic GαqGTP inhibitor perturbs the downstream signaling of PLCβ3. **(A)** Modeled structure of HTH tethered iLID and the cartoon depicting the inhibitor design. HTH: pink, iLID: gray, Jα helix of iLID: light green, FMN: blue. **(B)** Activation of GRPR in HeLa cells results in an uninterrupted PIP2 hydrolysis under no optical command in cells expressing Venus-iLID-HTH, Lyn-SspB, and mCherry-PH. **(C)** In a similar experiment, Venus-iLID-HTH was recruited to the membrane using blue light, and then bombesin was added. The cells didn’t exhibit a detectable PIP2 hydrolysis. **(D)** The plot shows the cytosolic PIP2 sensor dynamics with and without the optical command (n =20 for each condition). **(E)** Upon activation of GRPR, cells expressing Venus-iLID-HTH, Lyn-SspB, mCh-PH and Gαq showed a weakly attenuating PIP2 hydrolysis. Venus-iLID-HTH recruitment to the plasma membrane induced PIP2 hydrolysis attenuation and the blue light termination induced the PIP2 re-hydrolysis. The plot shows the PIP2 sensor dynamics in the cytosol with respect to iLID-HTH. **(F)** Upon expressing GRPR, Venus-iLID-HTH, Lyn-SspB and mCherry-DBD, GRPR activation in HeLa cells induced a robust mCherry-DBD translocation to the plasma membrane (white arrow) under no optical command. However, significant mCherry-DBD translocation was not observed while Venus-iLID-HTH was recruited to the plasma membrane using blue light (n =22 for the experiment and n=18 for the control). White arrow: mCh-DBD recruitment to the plasma membrane of the cells that are not exposed to blue light. The plot shows the dynamics of DAG formation. **(G)** HeLa cells expressing GRPR, Venus-iLID-HTH, and Lyn-SspB were treated with 2.28 μM Fluo-4 AM for 30 min at room temperature. GRPR was activated with 1 μM bombesin, and the increase in the Fluo-4 AM fluorescence intensity is examined. Minor fluorescence increase was observed in cells where Venus-iLID-HTH is recruited to the plasma membrane upon blue light. However, a significant calcium release was observed with the highly increased Fluo-4 AM fluorescence intensity in cells that are not subjected to the optical command (n=20 for each condition). Yellow arrow: minor Fluo-4 AM fluorescence increase in cells exposed to blue light upon GRPR activation. **(H)** The whisker box plot shows the relative Fluo-4 AM fluorescence increase upon activation, compared to the basal level of Fluo-4 AM fluorescence in cells with plasma membrane recruited Venus-iLID-HTH and cytosolic Venus-iLID-HTH. Average curves were plotted using cells from ≥3 independent experiments. The error bars represent SEM (standard error of the mean). The scale bar = 5 µm. mCh: mCherry; iLID: improved Light Induced Dimerization module; DBD: DAG Binding Domain. The blue box indicates the blue light exposure.

To examine the feasibility of optically switching ON and OFF of PLCβ signaling, we expressed GRPR, Venus-iLID-HTH, Lyn-SspB, and mCherry-PH in Gαq expressing HeLa cells. We have previously shown that overexpression of Gαq leads to weakly or non-attenuating PIP2 hydrolysis due to the elevated GαqGTP: PLCβ ratio and excessive Gβγ generation upon Gαq pathway activation^42^. In a cell with sustained PIP2 hydrolysis induced by GRPR activation (Fig. 2E- t= 0 s), mCherry-PH was rapidly recovered to the plasma membrane upon the blue light exposure induced Venus-iLID-HTH plasma membrane recruitment (Fig. 2E and the plot -blue box). Interestingly, the blue light termination caused a synchronized Venus and mCherry re-translocation to the cytosol (Fig. 2E- 300 s). The repeated blue light switching ON-OFF allowed repeated control of PIP2 hydrolysis, indicating the reversibility of iLID-HTH-induced PIP2 hydrolysis inhibition (Fig. 2E).

In addition to PIP2, we also show that the optogenetic control of PLCβ activity is reflected at the DAG-level. During time-lapse imaging of Venus and mCherry in cells expressing Venus-iLID-HTH, Lyn-SspB, and mCherry-DBD (DAG sensor),^43^ we exposed only selected cells in a field of vision to localized blue light (Fig.2F). Upon activation of GRPR, cells without blue light exposure showed DAG formation as indicated by the plasma membrane recruitment of the mCherry-DBD (Fig.2F- left, white arrow). On the contrary, blue light-exposed cells failed to show a visually detectable mCherry-DBD recruitment to the plasma membrane (Fig. 2F-right). The plot however shows a minor reduction in mCherry fluorescence in the cytosol, indicating only a minor DAG formation. Though cytosolic DAG sensor fluorescence reduction in blue light exposed cells is minor compared to the control cells, it is significantly different from the pre-GRPR activation level mCherry-DBD fluorescence at the cytosol (Fig. 2F-plot*, Fig.S3C, one-way ANOVA*: F*_3,63_ =383.72, *p* = <0.0001; Supplemental Table S6, A, and B).

We have observed that even residual PLCβ activity can induce sufficient PIP2 hydrolysis to produce detectable calcium mobilizations. Therefore, to examine the extent of PLCβ activity inhibition by iLID-HTH, we examined the Ca^2+^ responses with and without recruiting the HTH to the cell membrane in HeLa cells expressing mCherry-iLID-HTH and Lyn-SspB. We incubated cells for 30 min at room temperature with the fluorescent calcium indicator Fluo-4 AM (2.28 μM) before washing cells. Here, we imaged Fluo-4 AM using 515 nm excitation and 542 nm emission to prevent the iLID activation at unintended times. As expected, control cells activated by 1 μM of bombesin showed a sustained and significant Fluo-4 AM fluorescence increase (Fig. 2G-left), while the cells exposed to blue light in which mCherry-iLID-HTH recruited to the plasma membrane showed only a minor fluorescence increase, indicating a residual PLCβ activation (Fig.2G-right, yellow arrow). Similarly, normalized maximum Fluo-4 AM fluorescence show that, though minor, GRPR activation-induced Fluo-4 AM fluorescence increase in blue light exposed cells is significantly different from the pre-GRPR activation level Fluo-4 AM fluorescence intensity (Fig.2H) (one-way ANOVA*: F*_3,76_= 196.747, *p* = <0.0001; Supplemental Table S7, A, and B). This data agrees with both the observed minor PIP2 hydrolysis and DAG formation under optogenetic GαqGTP inhibition (Fig. 2C and 2F).

Therefore, to achieve complete optogenetic inhibition of GαqGTP-induced PLCβ signaling, we next engineered five inhibitor variants either by truncating (HTH 31-mer, HTH 27-mer, HTH 22-mer) or truncating and mutating (I860A in HTH 27-mer and P862H in HTH 27-mer) as described in Fig. 3A and table S1. Here the goal was to obtain a small peptide structure that can only be available upon optical command; however, completely unavailable to interact with GαqGTP in the absence of blue light. Since structural data show that Y847, I848, and P849 are not a part of the helix and D850 and D851 do not show strong interactions with GαqGTP, we incrementally truncated these residues to generate HTH 31-mer and 27-mer variants. Additionally, since L879, I880, G881, and E882 don’t exhibit strong interactions with GαqGTP, we also truncated those residues in HTH 27-mer variant. It has been shown that the extreme Ct domain of the HTH peptide (S867-E882) possesses an autoinhibitory effect by binding to a highly conserved cleft between the C2 domains of the catalytic core and TIM barrel near the active site of PLCβ3^44^. Therefore, we truncated that domain and generated the HTH 22-mer to examine if this domain influences HTH-GαqGTP interactions. We compared the extent of the PIP2 hydrolysis inhibition of each variant in HeLa cells additionally expressing GRPR, mCherry-PH, and Lyn-SspB. We first exposed cells to blue light (Fig. 3B-at 50 sec) and then to bombesin (Fig. 3B- at 120 sec). PIP2 hydrolysis inhibition by Venus-iLID-HTH 22-mer and the HTH 27-mer with P862H mutation was compromised compared to the other HTH variants (Fig. 3B- orange and green curves). We propose that HTH 22-mer showed only a partial inhibition, likely due to some extreme Ct residues also interacting with GαqGTP. The HTH 27-mer with P862H failed as an inhibitor since P862 may be required for the HTH secondary structure formation^45,17^. Compared to the inhibition observed with the 31-mer and 36-mer, the 27-mer HTH produced an enhanced inhibition (Fig. 3B-dark brown curve). However, the maximum PIP2 hydrolysis inhibition was observed with the HTH 27-mer with I860A mutation (Fig. 3B- black curve). This is inconsistent with the reported enhanced lipase activity of PLCβ3-I860A^17^. One-way ANOVA showed that the maximum PIP2 hydrolysis in Venus-iLID-I860A-HTH 27-mer significantly differs from the Venus-ilID-HTH that carries the entire 36-mer HTH (Fig. 3C, one-way ANOVA: *F_1, 38_* = 323.27216, *p* = 0; Table S8, A, and B). This data clearly indicated that iLID-HTH (I860A) can completely disrupt GαqGTP-PLCβ signaling. Therefore, we decided to employ the I860A-HTH 27-mer, which we named dHTH, to engineer the optogenetic PLCβ inhibitor further.

**Figure. 3:**
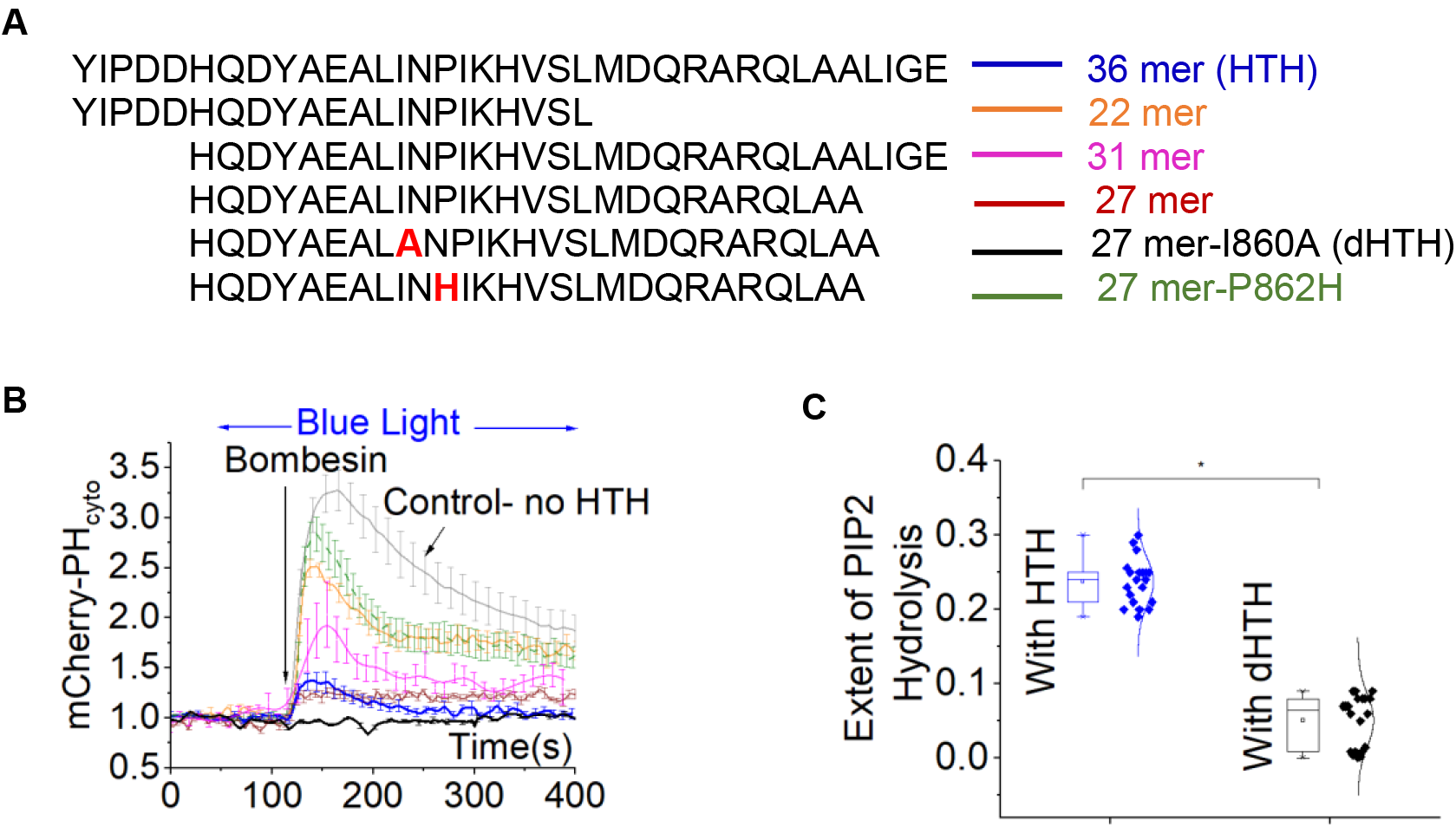
Optimization of iLID-based optogenetic GαqGTP. **(A)** Amino acid sequences alignment of Venus-ilID-HTH variants. **(B)** Comparison of the extent of PIP2 hydrolysis with Venus-iLID-HTH variants in HeLa cells expressing GRPR, Lyn-SspB, and mCherry-PH (n=15 for each variant and n= 20 for control). **(C)** The whisker box plot shows the extents of PIP2 hydrolysis in HeLa cells expressing GRPR, mCherry-PH, Lyn-SspB, and Venus-iLID-HTH or Venus-iLID-dHTH (n =20 for each variant). Average curves were plotted using cells from ≥3 independent experiments. The error bars represent SEM (standard error of the mean).

### 2.3 Optimization of optogenetic inhibitor for subcellular signaling

To examine the subcellular signaling inhibition, we recruited mCherry-iLID-dHTH to one side of a cell in which Gαq GPCRs are already activated, and PIP2 is hydrolyzed. Though mCherry accumulated more at the blue light exposed site (Fig. 4A, white arrow), it also showed a significant accumulation at distal plasma membrane regions. More importantly, the cell exhibited a global PIP2 recovery (Fig. 4A, yellow arrow). Though the blue light is localized, the activated iLID in the adjacent cytosol appears to be diffusing globally, inducing a cell-wide inhibition. Unfortunately, this global GαqGTP-PLCβ inhibition upon localized optical activation reduces the utility of iLID-dHTH in subcellular signaling control.

**Figure. 4:**
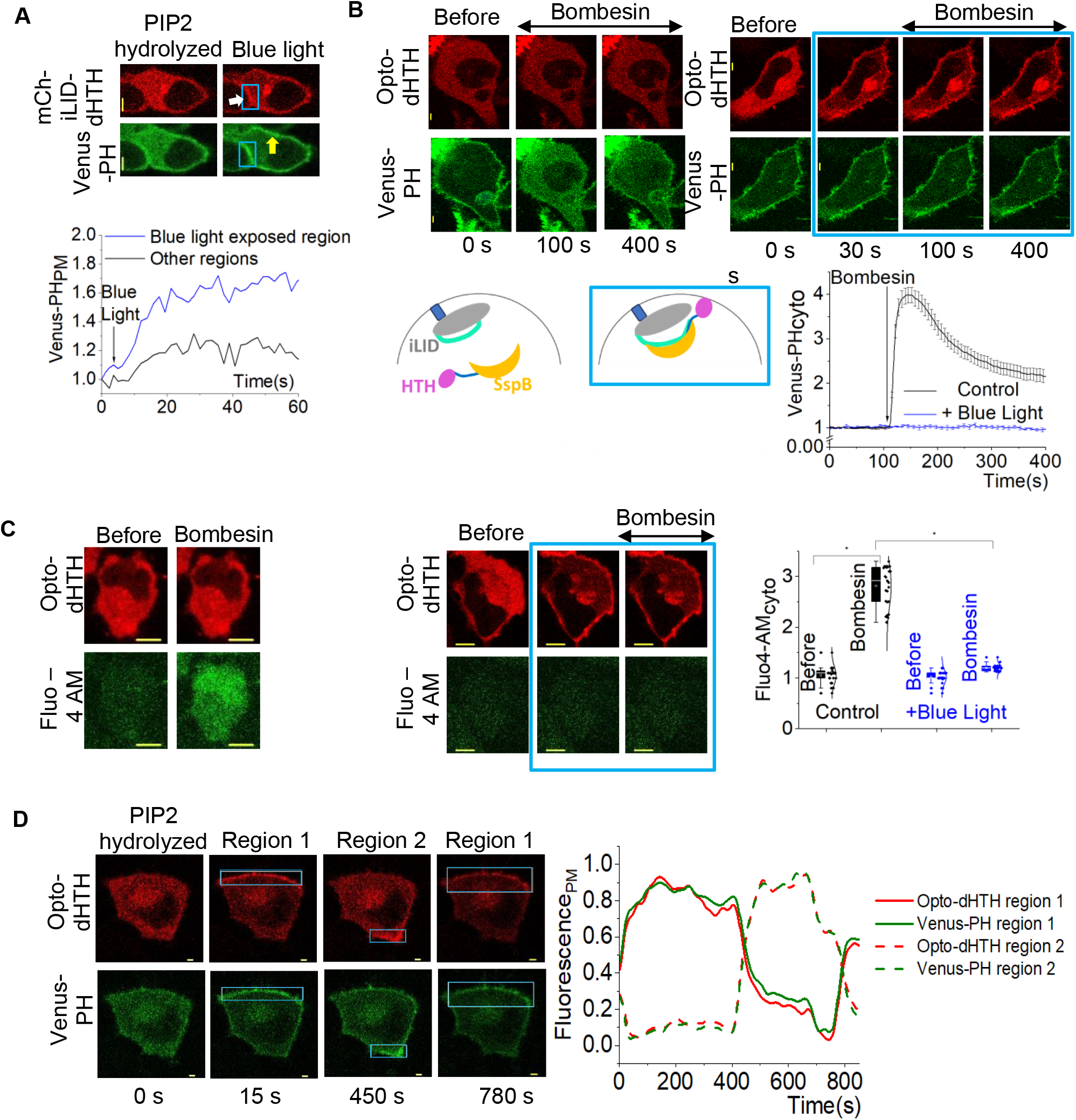
Engineering and validation of optogenetic GαqGTP inhibitor to control subcellular signaling. **(A)** Cell images show although the mCh-iLID-dHTH recruitment was significantly localized, the PIP2 hydrolysis was globally inhibited. White arrow: mCh-iLID-dHTH recruitment to the blue light exposed regions of the cell. Yellow arrow: PIP2 hydrolysis inhibition in unintended areas of the cell. The plot shows the plasma membrane recruitment of the PIP2 sensor in the blue light-exposed region and an unintended area of the cell. **(B)** Upon addition of 1 μM bombesin, HeLa cells expressing GRPR, Venus-PH, iLID-CAAX, and Opto-dHTH didn’t exhibit PIP2 hydrolysis with the optical command indicating its inhibitory effect. The plot shows the cytosolic PIP2 sensor dynamics with and without the optical command (n =25 for each condition). **(C)** HeLa cells expressing GRPR, Opto-dHTH, and iLID-CAAX were incubated with 2.28 μM of Fluo-4 AM. The increase in the Fluo-4 fluorescence intensity was examined upon GRPR activation. A detectable fluorescence increase was not observed in cells with plasma membrane recruited Opto-dHTH, while the Fluo-4 AM fluorescence significantly increased in the cells with cytosolic Opto-dHTH (n =30 for each condition). The whisker box plot shows the relative Fluo-4 AM fluorescence increase upon activation, compared to the basal level of Fluo-4 AM fluorescence in cells with plasma membrane recruited Opto-dHTH and cytosolic Opto-dHTH. **(D)** Subcellular inhibition of GαqGTP-induced PIP2 hydrolysis upon recruitment of Opto-dHTH into a confined membrane region. The blue box indicates the localized blue light. The plot shows the PIP2 sensor dynamics at the plasma membrane. Average curves were plotted using cells from ≥3 independent experiments. The error bars represent SEM (standard error of the mean). The scale bar = 5 µm.

To prevent the diffusion of activated iLID to unintended regions, we next anchored iLID to the plasma membrane (iLID-CAAX) while making dHTH-tethered SspB cytosolic (dHTH-mCherry-SspB). We generated three inhibitor variants to optimize spatial freedom for dHTH and enhance GαqGTP interactions with dHTH (Fig. S4A). In variant 1, dHTH was directly linked to mCherry (Fig.S4Ai), while in variant 2, we placed a 51-residue linker resembling the hypervariable region of PLCβ3 (*Ser* 882 to S*er* 933) (Fig. S4Aii) between dHTH and mCherry. This region links the proximal C terminal domain (CTD) and distal CTD of PLCβ3 (PDB ID: 4GNK). In variant 3, we used a truncated 17 residue linker (*Ser* 882 to *Gln* 899) from the same hypervariable region (Fig. S4Aiii). Upon expression in HeLa cells together with iLID-CAAX, all three variants could be locally recruited to the plasma membrane (Fig.S4A-cell images). We next examined the ability of variants1-3 to inhibit PLCβ signaling, first by globally recruiting them to the plasma membrane in HeLa cells expressing GRPR, iLID-CAAX, and Venus-PH, and then activating GRPR (Fig. S4B-C and Fig.4B). Regardless of the blue light exposure, we observed a significant PIP2 hydrolysis upon GRPR activation indicating that variant 1 only provided a minor inhibition of PIP2 hydrolysis (Fig. S4B). Though greater than variant 1, variant 2 (with 51 residue linker) also provided a partial PLCβ inhibition (Fig. S4C). The data collectively suggested though dHTH is a highly efficient inhibitor, in variant 1, it is spatially constrained to interact with GαqGTP, while in variant2, the 51-residue linker is excessively flexible, reducing dHTH reaction cross section with GαqGTP. This is somewhat analogous to having dHTH in the cytosol, which showed no detectable PLCβ inhibition. Interestingly, variant 3 with its 17 residue linker showed complete inhibition of GRPR-induced PIP2 hydrolysis (Fig. 4B). We named this inhibitor Opto-dHTH. Plasma membrane-recruited Opto-dHTH also inhibited DAG formation (Fig. S4D). The Alphafold2 modeled structure of Opto-dHTH further indicates that dHTH has a favorable orientation to interact with GαqGTP on the plasma membrane, while its SspB interacts with the membrane-bound iLID (Fig. S4E). We next examined whether Opto-dHTH can induce complete PLCβ inhibition using GRPR activation-induced calcium mobilization in cells with and without blue light exposure (Fig. 4C). Cells without blue light exposure produced a robust increase in cytosolic calcium (Fig. 4C-left). However, the mean Fluo-4 AM fluorescence intensities before and after GRPR activation in cells exposed to blue light were not significantly different, indicating the highly efficient inhibition of PLCβ signaling by Opto-dHTH (Fig. 4C-plot), (one-way ANOVA: *F_3, 96_* = 369.159, *p* = <0.0001; Supplemental Table S9A, B). The data suggests that Opto-dHTH can inhibit even the residual levels of PLCβ signaling.

To examine the feasibility of subcellular GαqGTP-PLCβ interaction inhibitions, we expressed GRPR, Opto-dHTH, iLID-CAAX, and Venus-PH in HeLa cells. In a cell where the PIP2 is hydrolyzed upon GRPR activation, we exposed a confined plasma membrane region to blue light and recruited Opto-dHTH locally (Fig. 4D- blue boxes). Indicating the localized inhibition of GαqGTP-PLCβ interactions, the PIP2 sensor returned to the blue light-exposed area, suggesting the localized PIP2 generation. Switching the blue light exposed area of the same cell shows the user-defined reversible spatiotemporal control of PLCβ activity (Fig. 4D and supplemental Movies S1 and S2). The localized inhibition was reversible and ceased upon blue light termination. This data indicated that not only Opto-dHTH is an efficient inhibitor of PLCβ signaling, but it also possesses the ability to control subcellular signaling in a temporally precise manner.

### 2.4. Opto-dHTH does not disrupt Gβγ signaling or GαqGTP-GRK2 interactions

To examine whether Opto-dHTH interferes with GPCR activation-induced G protein heterotrimer dissociation and free Gβγ generation, we employed Gγ9 translocation assay^46, 47^. Gγ9 translocates from the plasma membrane to internal membranes upon GPCR activation and has been developed as an assay to probe for cellular and subcellular GPCR-G protein activities^46, 48–50^. Here, we examined whether GRPR activation-induced free Gβγ generation is intact in cells in the presence of Opto-dHTH-mediated GαqGTP-PLCβ interactions inhibition at the plasma membrane. We expressed GRPR, Opto-dHTH, iLID-CAAX, mCherry-Gγ9, and Gαq in HeLa cells (Fig. 5A). We have previously shown that the Gβγ translocation upon Gαq-GPCR activation is only visible in HeLa cells upon Gαq expression because of the limited endogenous Gαq availability^42^. The baseline normalized endomembrane Gγ9 fluorescence shows that Opto-dHTH activation does not interfere with GRPR activation-induced free Gβγ generation and subsequent translocation (one-way ANOVA: *F_1, 33_* = 0.1326, *p* = 0.71804; Supplemental Table S10A-B, Fig. 5A, and S5A). In addition to Gγ9 translocation, we examined the Gβγ−activated phosphoinositide-3-kinases (PI3Ks) induced PIP3 generation in RAW264.7 cells. We employed RAW264.7 cells because, as shown previously, HeLa cells did not show an observable PIP3 generation due to the lack of high membrane affinity Ggs^51, 52^. PIP3 generation was measured using the translocation of the fluorescently tagged PIP3 sensor (Akt-PH-Venus) from cytosol to the plasma membrane. Since GRPR-induced PIP3 generation in cells only with endogenous Gαq is minor, and is enhanced by Gαq expression (Fig. 5B-left), for this experiment, we use RAW264.7 cells expressing GRPR, Opto-dHTH, iLID-CAAX, Akt-PH-Venus, and Gαq. GRPR was activated after recruiting Opto-dHTH to the plasma membrane (Fig. 5B-middle). Similar to the control RAW264.7 cells without the Opto-dHTH expression, the cells with the membrane recruited Opto-dHTH showed a detectable PIP3 generation (one-way ANOVA: *F_1, 18_* = 0.42163, *p* = 0.5243; Supplemental Table S11A-B, Fig. 5B and Fig. S5B). This data collectively suggests that Opto-dHTH does not have a significant influence on heterotrimer activation or Gβγ signaling.

**Figure. 5:**
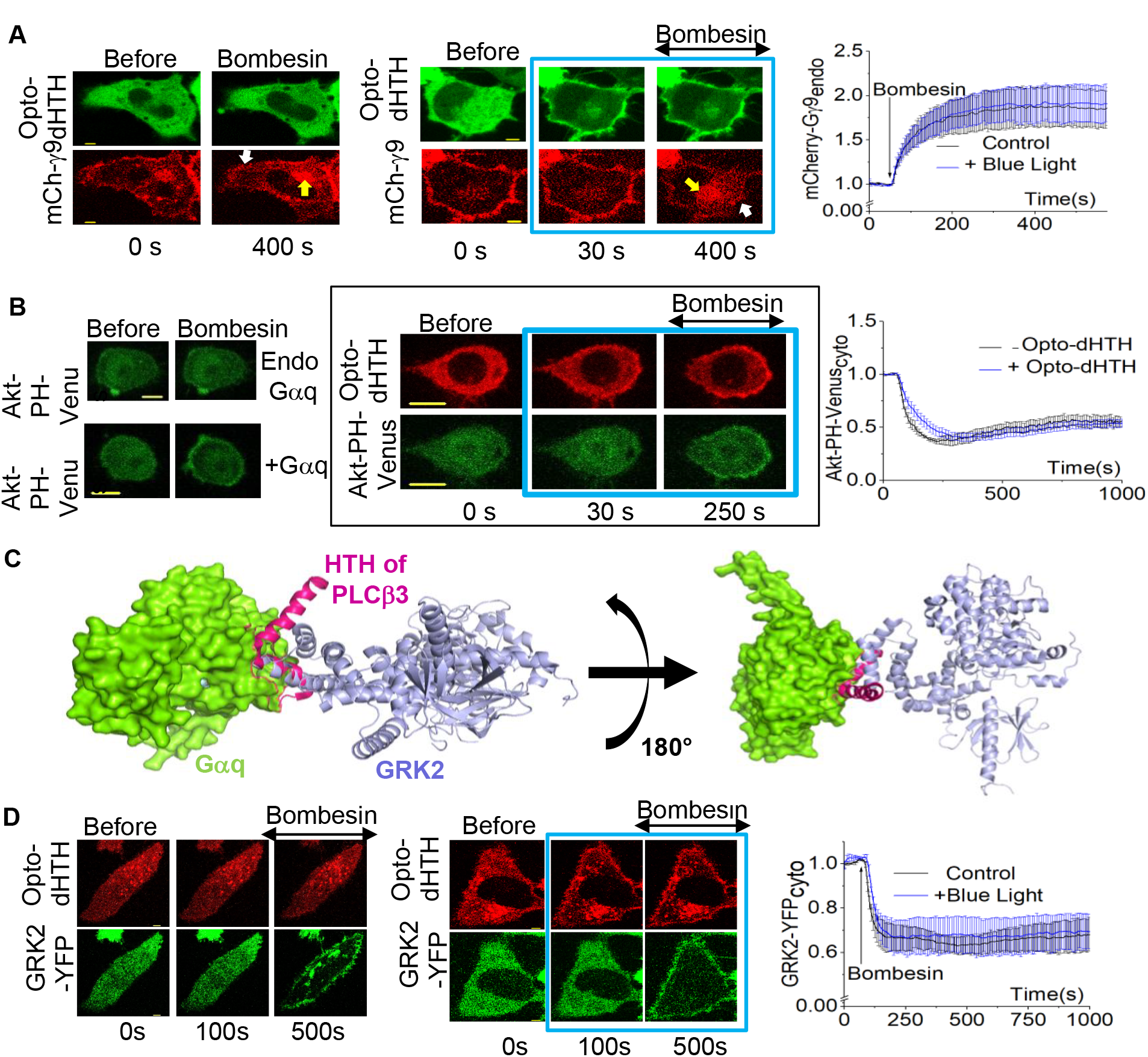
dHTH selectively inhibits GαqGTP without interrupting the dissociated Gβγ-induced signaling and GRK2 binding to GαqGTP. **(A)** HeLa cells expressing GRPR, Gαq-CFP, mCh-Gγ9, iLID-CAAX, and Opto-dHTH were activated with 1 μM bombesin. Robust Gγ9 translocation was observed in the cells with and without the recruitment of Opto-dHTH to the plasma membrane (n =30 for each condition). The plot shows the baseline normalized mCh fluorescence in endomembranes over time. The loss of Gγ9 from the plasma membrane and accumulation of Gγ9 in internal membranes are indicated, respectively, by white and yellow arrows. **(B)** GRPR and Akt-PH-Venus were expressed in RAW 264.7 cells. A detectable PIP3 generation was not observed with the endogenous Gαq in RAW 264.7 cells (Fig. 5B-left-top). GRPR, Gαq-CFP, and Akt-PH-Venus were expressed in RAW 264.7 cells, and a significant PIP3 generation was observed (Fig. 5B-left-bottom). GRPR, Gαq-CFP, Opto-dHTH, iLID-CAAX, and Akt-PH-Venus were expressed in RAW 264.7 cells. Opto-dHTH was recruited to the plasma membrane and GRPR was activated as described earlier. A significant Akt-PH-Venus recruitment to the plasma membrane was observed, indicating similar PIP3 generation extent to the control cells lacking Opto-dHTH expression. The corresponding plot shows PIP3 sensor dynamics in the cytosol of the cells. **(C)** Overlapping crystal structures of Gαq-PLCβ3 (PDB ID: 4GNK) and Gαq-GRK2 (PDB ID: 2BCJ) showing the interactions of the HTH domain and GRK2 with Gαq. Gαq: green; HTH: pink; GRK2: light purple. **(D)** GRPR, Gαq-CFP, GRK2-YFP, iLID-CAAX, and Opto-dHTH were expressed in HeLa cells. Upon activation of GRPR with bombesin (1μM), GRK2-YFP was recruited to the cell membranes in which Opto-dHTH was on the membrane and in the cytosol (n = 27 for the condition with blue light and n= 16 for the condition without blue light). The plot shows the dynamics of cytosolic GRK2-YFP. Average curves were plotted using cells from ≥3 independent experiments. The error bars represent SEM (standard error of the mean). The scale bar = 5 µm. GRK2: G protein-coupled receptor kinase 2; Gβγ: G protein βγ. The blue box represents the blue light exposure.

G protein-coupled receptor kinases (GRKs) desensitize GPCRs and prevent hyperactivation of downstream signaling^53, 54,55^. Interestingly both GRK2 and the HTH domain of PLCβ interact with the switch region II of GαqGTP (Fig. 5C)^56^. Although they don’t interact through the same residues, next, we examined whether the Opto-dHTH-Gαq interactions perturb GRK2 interactions with Gαq. Using HeLa cells expressing GRPR, Opto-dHTH, iLID-CAAX, GRK2-YFP, and Gaq, we examined the GRK2-YFP recruitment to the plasma membrane, both with and without blue light exposure (Fig. 5D). Regardless of the blue light, cells showed similar and robust GRK2-YFP recruitments to the plasma membrane (one-way ANOVA: *F_1, 28_* = 0.8247, *p* = 0.37155; Supplemental Table S12A-B, Fig. 5D, Fig. S5C). This clearly indicates that GRK2-GαqGTP interactions are not affected by GαqGTP – dHTH interactions.

### 2.5 Subcellular optogenetic control of GαqGTP-PLCβ signaling

Directional cell migration is crucial in many physiological and pathological processes, such as immune system function, tissue remodeling, and embryonic development^57, 58^. GNAQ that encodes for Gαq has been identified as a cancer driver gene implicated in uveal melanoma, lung adenocarcinoma, and cutaneous melanoma^42, 59–62^. However, when a migratory cell senses a chemokine gradient that activates the Gαq-GPCR pathway, it is unclear how cells reorient and respond because methods to produce spatially confined GαqGTP activity in a subcellular region of a single cell are lacking. We previously showed that Gβγ regulates the leading edge cytoskeleton remodeling and directional cell migration of macrophages using subcellular activation of blue opsin photoreceptor, a Gi/o GPCR^57^. On the contrary, Gαq has been shown to involve cytoskeletal retraction through Rho A activation^63, 64^. Since Gαq-GPCR activation induces both GαqGTP and Gβγ signaling, and they have opposing effects, we examined how the two transducers collectively regulate Gαq-GPCR-induced directional cell migration.

We first examined how macrophages adhered to cell culture dishes (Fig. 6A- first image) respond to the asymmetric disruption of Gαq-PLCβ interactions across the cells using Opto-dHTH. In GRPR, iLID-CAAX, and Opto-dHTH expressing RAW264.7 mouse macrophage cells on imaging glass-bottomed dishes, we first activated GRPR by exposing cells to bombesin, and allowing for PIP2 to be hydrolyzed. Using localized blue light, we recruited Opto-dHTH to a confined membrane region of the cell (Fig. 6A, 2 and 8 min, Movie S3). Interestingly, the cell started migrating away from the blue light-exposed region (Fig. 6A, 8 min). When we switched the localized blue light to the opposite side, Opto-dHTH, too, switched the localization, and the cell started migrating away from the blue light again (Fig. 6A, 10 min).

**Figure. 6:**
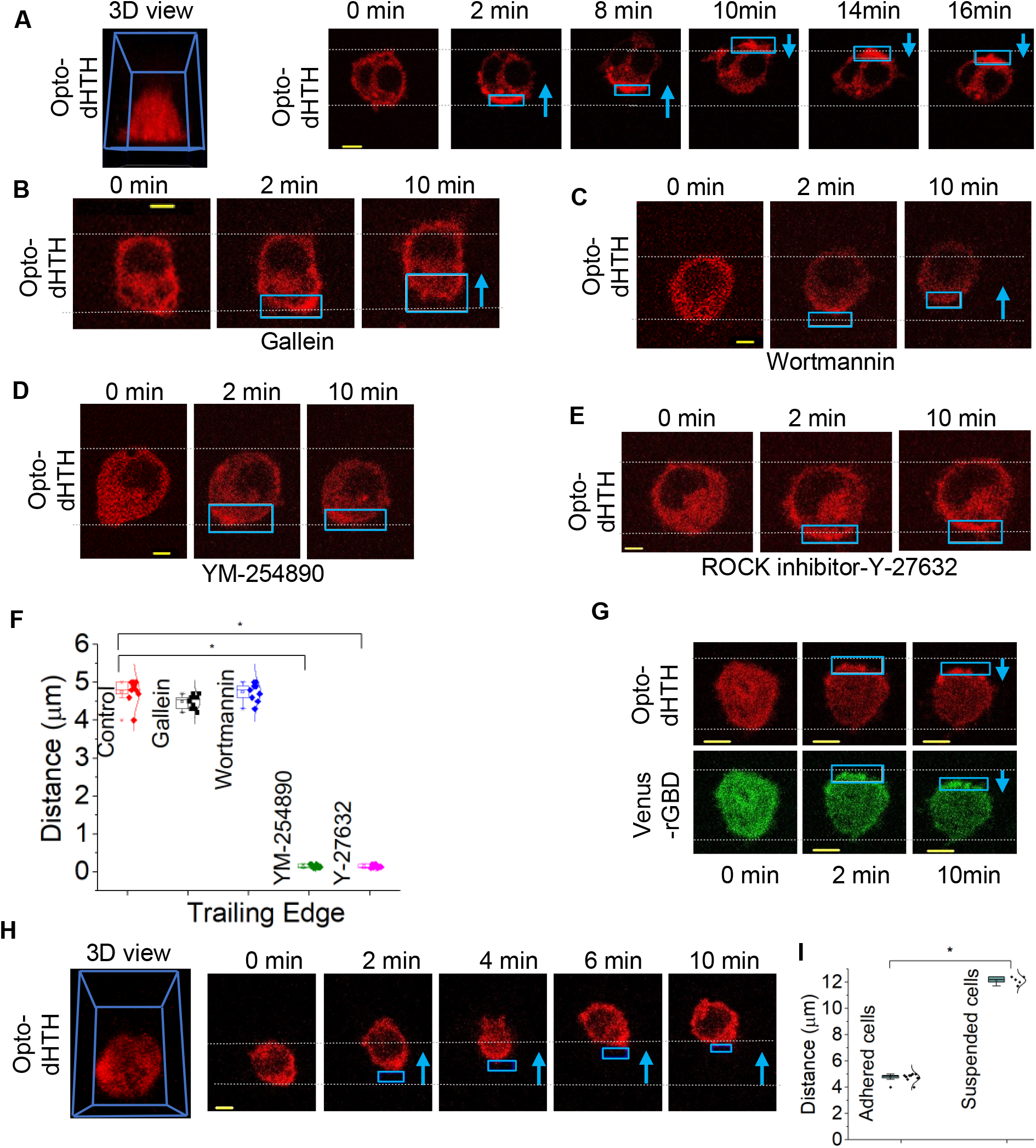
Subcellular recruitment of Opto-dHTH induces migration of Gq-GPCR activated RAW264.7 cells. **(A)** 3D view of a RAW264.7 cell adhered to the imaging glass bottomed dish is shown in the first cell image. RAW264.7 cells expressing GRPR, iLID-CAAX, and Opto-dHTH were activated with 1 μM of bombesin. Then Opto-dHTH was recruited to one side of the cell using localized blue light (*blue box*) and the cell started migrating opposite to the optical command (n =10). Blue arrow indicates the direction of cell migration. **(B-F)** Similar experiments were performed in cells treated with pharmacological inhibitors for Gβγ, PI3K, GαqGTP, and ROCK (Rho-associated protein kinase). **(B)** RAW264.7 cells treated with the Gβγ inhibitor, gallein (10 μm, 30 min, 37 °C) also migrated opposite to blue light (n =10). **(C)** RAW264.7 cells treated with 50nM wortmannin also showed similar migration opposite to blue light (n =9). **(D)** RAW264.7 cells treated with the GαqGTP inhibitor, YM-254890 (500 nM) didn’t show a detectable migration (n =9). **(E)** RAW264.7 cells treated with the ROCK inhibitor, Y-27632 (10 μM) didn’t exhibit a detectable migration (n =9). **(F)** The whisker box plot shows the extent of movement of the peripheries of the trailing edge of RAW264.7 cells described in A–E, within 10 minutes. **(G)** GRPR, Opto-dHTH, iLID-CAAX, and RhoA sensor, Venus-rGBD were expressed in RAW264.7 cells. Opto-dHTH was recruited to one side of the cell after GRPR activation. Venus-rGBD was accumulated on the Opto-dHTH recruited region of the cells. (n =5). (**H)** 3D view of a RAW264.7 cell suspended in the iso-dense media is shown in the first cell image. RAW264.7 cells expressing GRPR, iLID-CAAX, and Opto-dHTH were suspended in 23% ficoll and activated with 1 mM of bombesin. Suspended cells showed a higher migration than the adherent cells, upon Opto-dHTH recruitment to one side of the cell (n= 5). **(I)** The whisker plot shows the Opto-dHTH induced migration distances of suspended and adherent RAW264.7 cells within 10 minutes (n= 10 for adherent cells and n= 5 for suspended cells).Average curves were plotted using cells from ≥3 independent experiments. The error bars represent SEM (standard error of mean). The scale bar = 5 µm. PI3K: phosphatidylinositol-3 kinase; rGBD: Rhotekin G protein binding domain.

To understand the molecular reasoning underlying this migratory behavior, we performed the experiment under various pharmacological perturbations, including 10µM gallein (Gβγ inhibitor), 50 nM wortmannin (PI3K or phosphatidylinositol-3 kinase inhibitor), 10 µM Y-27632 (Rho kinase inhibitor), and 500 nM YM-254890 (Gαq inhibitor) (Fig.6B-F)^57^. Both gallein and wortmannin-treated cells exhibited migrations similar to the control (Fig.6B, C and F). Gβγ-PI3K signaling plays a prominent role in Gi/o -GPCR induced cell migration when their signaling is locally activated^57, 65^. Although Opto-dHTH allows for GαqGTP-PLCβ signaling to be confined to one side of the cell, it does not influence Gβγ signaling (Fig. 5B), creating a uniform Gβγ signaling paradigm across the cell. Therefore, it is not surprising that neither gallein nor wortmannin influenced the observed migration. As we described above, limited Gβγ and subsequent PI3K activity in these cells upon Gq-pathway activation could be another reason for the lack of influence by gallein and wortmannin. Cells exposed to 500 nM YM-254890 did not show a detectable directional migration (Fig. 6D). This is not unexpected since YM-254890 is a Gαq-specific inhibitor^15^, which inhibits GDP to GTP exchange on Gαq^66^. RAW264.7 cells exposed to 10 μM of the Rho-kinase inhibitor, Y-27632 (30 min at 37°C)^67^ didn’t show Opto-dHTH-induced directional migration away from blue light (Fig. 6E). This can be understood because RhoA is a key regulator of collective cell migration^68^, and RhoA causes myosin light chain contractions upon activation^69, 70^. The plot shows that, compared to the control, as well as gallein and wortmannin exposed cells, trailing edge retractions in cells exposed to Y-27632 and YM-254890 were significantly decreased (Fig. 6F) (one-way ANOVA: *F_4, 41_* = 1446.833, *p* = <0.0001; Table S14A and B). To further confirm the effect of Opto-dHTH on RhoA activation, we expressed Venus-rGBD^71^, a RhoA sensor in RAW264.7 cells expressing GRPR, Opto-dHTH, iLID-CAAX. The Venus-rGBD recruitment to the plasma membrane has been used as an indicator for RhoA activation in cells^71^. Opto-dHTH was recruited to one side of the GRPR already activated cell, and detectable synchronized recruitment of Venus-rGBD to the blue light-exposed side of the cell was observed (Fig. 6G). This indicated that the recruitment of Opto-dHTH enhances RhoA activation on the Opto-dHTH recruited side. As a control, compared to Lyn-mRFP expressing cells (FigS6A), Lyn-mRFP-dHTH cells shows better RhoGEFs recruitment to the plasma membrane upon GRPR activation (Fig. S6B-white arrow). This data suggests that when dHTH is bound to GαqGTP, although PLCβ cannot, RhoGEFs can interact with it, and induces a robust RhoA activation. On the contrary, it is likely that in the presence of PLCβ, RhoGEFs interactions with GαqGTP is suppressed.

To compare the localized Gαq-GPCR activation-induced cell migration with and without the influence of dHTH, we expressed melanopsin, emiRFP670 and Lyn-mRFP or Lyn-mRFP-dHTH. Melanopsin is a photoreceptor GPCR found in intrinsically photoreceptive retinal ganglion cells-ipRGCs from the retina that activated both Gq and Gi/o pathways with near equal efficiencies^72, 73^. We activated melanopsin in a localized region of a cell using blue light while imaging emiRFP670 using 637 nm excitation to image cell migration without globally activating the receptor (Fig.S6D). To eliminate Gi/o signaling influence, this experiment was conducted in the presence of 0.05 μg/ml pertussis toxin^74^. Upon melanopsin activation, Lyn-mRFP expressing RAW264.7 cells showed a minor migration away from the blue light (Fig.S6D- top raw). The cells expressing Lyn-mRFP-dHTH showed significantly higher migration than the migration observed with Lyn-mRFP (Fig. S6E) (one-way ANOVA: *F_1, 10_* = 14.0369, *p* = 0.00381; Table S15A and B). When localized melanopsin activates Gαq-heterotrimers in the absence of Lyn-mRFP-dHTH, we propose that PLCβ compete for the generated GαqGTP, suppressing RhoGEF and RhoA activation and, subsequently, actomyosin contractility.

We next compared this asymmetric RhoA activation-triggered macrophage migration with the localized blue opsin activation-induced macrophage migration, which shows prominent lamellipodia formation at the opsin-activated leading edge. Upon blue opsin activation, RAW264.7 cells expressing Akt-PH-mCherry (a PIP3 sensor that translocates to the plasma membrane from the cytosol) showed a robust PIP3 generation and lamellipodia formation at the blue light-exposed leading edge of the cell (Fig. S6F). We have extensively characterized this blue opsin-induced, Gi/o-Gβγ mediated migration characterized by the prominent leading-edge activity starting before the trailing-edge retraction^57^. On the contrary, not only the RhoA-induced migration first showed retraction of the trailing edge, but it was also accompanied by relatively slow migration of the cells without prominent lamellipodia formation at the leading edge (Fig. 6A). Careful examination indicated that, this proposed RhoA-based migration occurred under subcellular optogenetic inhibition of GαqGTP-PLCβ interactions bears the characteristic of the previously documented amoeboid cell migration^71^. Using an optogenetic RhoGEF suggested that upon RhoA activation, the cell membrane flows from leading to trailing edge, generating tangential viscous forces at the cell-liquid interface^71^. These forces drive the cells forward more swiftly in viscous media with low Reynolds numbers, following theoretical predictions made over a four-decades ago in an article titled “Life at low Reynolds number” ^75^. We, therefore, examined whether Opto-dHTH-directed migration in Gq-GPCR background recapitulates this “swimming-like” migration in a viscous medium (Fig. 6H). To prevent cell adhesion and provide the viscous medium, we suspended RAW264.7 cells expressing GRPR, Opto-dHTH, and iLID-CAAX in an iso-dense medium comprising 23% Ficoll containing 1 mM bombesin, using a modified protocol described previously^71^. Fig. 6H, the first image shows the spherical cell suspended in the isodense medium. Using the transparent nature of Ficoll, we recruited Opto-dHTH to one side of the cells using localized blue light, similar to adhered cells, suspended cells showed migration away from blue light, however, more aggressively (Fig. 6I) (one-way ANOVA: *F_1, 13_* = 2081.375, *p* = <0.0001; Table S13A and B).

Data presented collectively indicated that PLCβ-GαqGTP interactions suppress RhoGEF activation by GαqGTP, while dHTH-GαqGTP interactions do not. Therefore, Opto-dHTH recruitment to one side of the cell creates a RhoGEFs activity gradient, low on the opposite side and high on the recruited side. The sensitivity of Opto-dHTH-induced migration to ROCK inhibitor further indicates that the enhanced RhoGEFs and subsequent active RhoA at the Opto-dHTH side are the driving forces of the migration. We propose that locally activated Rho Kinase (ROCK) at the Opto-dHTH side reduces myosin light chain phosphatase (MLCP) activity, thus increasing phosphorylated MLCP and, subsequently, myosin light chain (MLC) phosphorylation, resulting in actomyosin contractility, creating the rear-end of the cell, as we described previously^57^. This mechanism is consistent with the previous findings that cells migrate away from the side of the cells where an optogenetic RhoGEF is localized^71^.

Data from Opto-dHTH under Gq-activated background, as well as Lyn-dHTH in melanopsin experiments, dually supported our hypothesis that PLCβ acts against GαqGTP-RhoGEFs interactions and retarding migration of adhered cells and cells experiencing chemokine gradients in 3D viscous fluidic media. The presented sample cell migration investigation using Opto-dHTH provided insights into the Gq-GPCR governed amoeboid-types cell migration that has not been sufficiently explored. Our findings helped build a likely mechanism that links Gq-GPCR, GNAQ, PLCβ and downstream signaling components underlying Gq-GPCR-induced cell migration (Fig. 7).

**Figure. 7:**
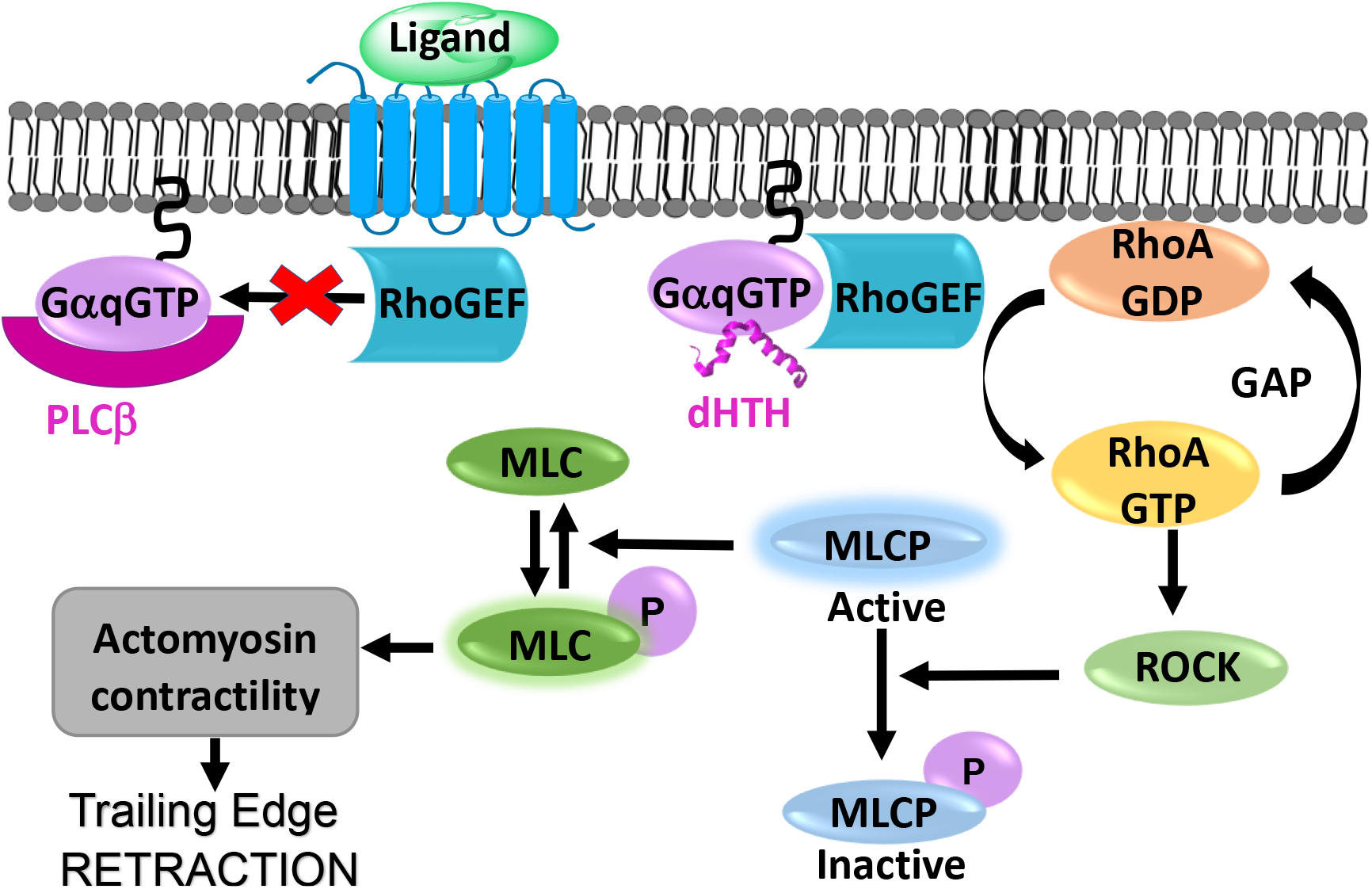
Proposed pathway for GαqGTP-induced RAW264.7 cell migration. Activation of Gαq-coupled GPCRs induces the heterotrimer dissociation, resulting in GαqGTP and free Gβγ. GαqGTP simultaneously induces the activation of multiple effectors such as PLCβ, RhoGEF, and GRK and, thus, several signaling pathways. Upon Gαq-coupled GPCR activation PLCβ and RhoGEFs compete for GαqGTP. When Opto-dHTH is bound to GαqGTP, more RhoGEFs interact with GαqGTP. Activated RhoGEFs induce trailing edge retraction by deactivating MLCP by activating the RhoA pathway. This mechanism increases phosphorylated MLC in the Opto-dHTH recruited side of the cell, promoting actomyosin contractility and subsequent retraction of the trailing edge.

### Conclusion

Our data demonstrate that Opto-dHTH can reversibly induce both cell-wide and subcellular inhibition of GαqGTP-PLCβ interactions and PLCβ signaling, while it neither perturbs Gαq heterotrimer activation, Gβγ generation, nor Gαq-GRK2 interactions. The data also indicated that GαqGTP-PLCβ interaction inhibition by Opto-dHTH is highly selective and efficient. Considering that even a few IP3 molecules upon PIP2 hydrolysis are sufficient to induce Ca^2+^ mobilization^76^, the observed complete lack of Ca^2+^ release indicates that Opto-dHTH induced PLCβ signaling inhibition is complete. These capabilities of Opto-dHTH can be attributed to its design and the small spatial volume occupancy at the GαqGTP surface. In the presence of globally activated Gαq-GPCRs, its ability to inhibit PLCβ signaling in subcellular regions and accompanied directional cell migration informs the molecular-level regulation of Gαq pathway-induced cell migration. The data also suggests that the migration directionality and extent are governed by the composition and the potency of competing interactions of effectors downstream of heterotrimer activation, including PI3Ks, RhoGEFs, RhoGTPases, and PLCβ. This is significant because compared to other Gα types, Gαq control many downstream effectors including PLCβ, and RhoGEFs, regulators of G protein signaling (RGS) 2 and 3, and GRK2, 3^77^. Therefore, it is not surprising that, besides cancer, the Gαq pathway is a major driver of many diseases, including vascular, cardiac, endocrine, and metabolic diseases^15, 78–80^. The abrupt and intense signaling and cell morphology changes induced by activated PLCβ usually make interrogation of other Gαq sub pathways challenging. The spatial and temporal control of PLCβ activity by Opto-dHTHs now allows for optically switching ON-OFF the PLCβ signaling from the overall Gαq-pathway response. Further, the lack of pharmacological inhibitors for PLCβ has been a long-standing obstacle in examining its signaling. Therefore, the presented work is exciting since we now show the feasibility of using membrane-targeted dHTH for constitutive permanent and Opto-dHTH for reversible, single-cell, and subcellular PLCβ activity inhibition. Finally, our data also shows a regulatory mechanism in which PLCβ significantly attenuates RhoGEFs signaling through its multipoint interactions with GαqGTP, also shedding light on to Gq-GPCR-RhoA-governed cell migration in viscous fluids. Further investigations are required to know if PLCβ similarly regulates other Gαq-effectors too. Collectively, the data inducates the potential significant utility of Opto-dHTH in probing Gq-PLCβ signaling regulation.

## 3. Materials and methods

### 3.1 Reagents

The reagents used were as follows: Bombesin (Tocris Bioscience), YM-254890 (Focus Biomolecules), wortmannin (Cayman Chemical), Gallein (TCI AMERICA), Pertussis toxin (Sigma Aldrich), Y-27632(Cayman Chemical), Ficoll400 (F8016; Sigma Aldrich), and 11-cis-retinal (National Eye Institute). According to the manufacturer’s instructions, all the reagents were dissolved in appropriate solvents and diluted in 1% Hank’s balanced salt solution supplemented with NaHCO_3_, or a regular cell culture medium, before adding to cells.

### 3.2 DNA constructs

DNA constructs used were as follows: GRPR, mRFP-HTH, mRFP, Lyn-mRFP-HTH, Lyn-mRFP, Lyn-mRFP-AsLOV2-HTH, CRY2-mCherry-HTH, Venus-iLID-HTH and variants, Lyn-SspB, mCherry-PH, Venus-PH, mCherry-DBD, mCherry-iLID-HTH, dHTH-mCherry-SspB, dHTH-51 residue linker-mCherry-SspB, Opto-dHTH, Gαq-CFP, GRK2-YFP, Venus-miniGq, CIBN-CAAX, iLID-CAAX (Addgene plasmid # 85680), melanopsin, Akt-PH-mCherry, Akt-PH-Venus, Venus-rGBD, and mCherry-Gγ9^11, 52, 72, 81^. GRPR was a kind gift from the laboratory of Dr. Zhou-Feng Chen at Washington University, St Louis, MO. mCherry-PH, Venus-PH, Gαq– CFP, mCherry-DBD, melanopsin, Akt-PH-mCherry, Akt-PH-Venus and fluorescently tagged Gγ9 were kindly provided by Dr. N. Gautam’s laboratory, Washington University in St Louis, MO. The plasmid GRK2-YFP was kindly provided by Dr. Moritz Bunemann, Department of Pharmacology and Clinical Pharmacy, Philipps-University Marburg, Germany. Venus-miniGq is a gift from Dr. Nevin Lambert, Augusta University, GA. All cloning was performed using Gibson assembly cloning (NEB). All cDNA constructs were confirmed by sequencing.

### 3.3 Cell culture and DNA transfection

RAW264.7 and HeLa cells were purchased from ATCC, USA. Recommended cell culture media; (RAW264.7 (RPMI/10%DFBS/1%PS), and HeLa (MEM/10%DFBS/1%PS) were used to subculture cells on 29 mm, 60 mm, or 100 mm cell culture dishes. For live-cell imaging experiments, cells were seeded on 29 mm glass-bottomed dishes at a density of 1 × 10^5^ cells. DNA transfections were performed using either Lipofectamine^®^ 2000 reagent (for HeLa cells) or electroporation (for RAW264.7 cells) according to the manufacturer’s recommended protocols. Briefly, for electroporation of RAW264.7 cells, the following method was used. Nucleofector solution (82 µL), Supplement solution (18 µL), and appropriate volumes of plasmid DNA for specific DNA combinations were mixed. In each electroporation experiment, ∼2-4 million cells were electroporated using the T-020 method of the Nucleofector™ 2b device (Lonza). Immediately after electroporation, cells were mixed with cell culture medium at 37 ⁰C and seeded on glass-bottomed dishes. Imaging was conducted after ∼5-6 h post-transfection, considering the high expression of constructs.

### 3.4 Live cell imaging, image analysis, and data processing

The methods, protocols, and parameters for live cell imaging are adapted from previously published work^43, 82, 83^. Briefly, live cell imaging experiments were performed using a spinning disk confocal imaging system (Andor Technology) with a 60X, 1.4 NA oil objective, and iXon ULTRA 897BVback-illuminated deep-cooled EMCCD camera. Photoactivation and Spatio-temporal light exposure on cells in regions of interest (ROI) was performed using a laser combiner with a 445 nm solid-state laser delivered using Andor® FRAP-PA (fluorescence recovery after photobleaching and photoactivation) unit in real-time, controlled by Andor iQ 3.1 software (Andor Technologies, Belfast, United Kingdom). Fluorescent proteins such as mCherry-PH, mCherry-γ9, mCherry -DBD, CRY2-mCherry-HTH, Lyn-mRFP-HTH, Lyn-mRFP-AsLOV2-HTH, Akt-PH-mCherry were imaged using 594 nm excitation−624 nm emission settings. Akt-PH-Venus, Venus-PH, Venus-iLID-HTH, Venus-miniGq, and GRK2-YFP were imaged using 515 nm excitation and 542 nm emission. mTurquoise-PH was imaged using 445 nm excitation and 478 nm emission, and Fluo-4 AM was imaged using 515 nm excitation and 542 nm emission, respectively. For global and confined optical activation of CRY2, LOV, and iLID expressing cells, 445 nm solid-state laser coupled to FRAP-PA was adjusted to deliver 145 nW power at the plane of cells, which scanned light illumination across the region of interest (ROI) at 1 ms/μm^2^. The time-lapse images were analyzed using Andor iQ 3.2 software by acquiring the mean pixel fluorescence intensity changes of the entire cell or selected area/regions of interest (ROIs). Briefly, the background intensity of images was subtracted from the intensities of the ROIs assigned to the desired areas of cells (plasma membrane, internal membranes, and cytosol) before intensity data collection from the time-lapse images. The intensity data from multiple cells were opened in Excel (Microsoft office®) and normalized to the baseline by dividing the whole data set by the average initial stable baseline value. Data were processed further using Origin-pro data analysis software (OriginLab®).

### 3.5 Structure determination of Opto-dHTH via AlphaFold

We used the amino acid sequences of SsrA, SspB, mCherry, HTH, and the corresponding linker to generate the modeled protein structure in Fig.S3E using AlphaFold2^84^. We fed these sequences to AlphaFold2 and built homology models. Structures that best fit the experimental structures were selected for protein folding validation. Finally, the protein preparation tool in Schrodinger Maestro13.3.121 was used to optimize the selected protein models. A similar procedure was used to generate the HTH-bound iLID structure in Fig. 2A and the structures of PLCβ isoforms used in Fig. S2B.

### 3.6 Experimental rigor and Statistical analysis

To eliminate potential biases or preconceived notions and improve the experimental rigor, we used the reagent-blinded-experimenter approach for the key findings of our study, i. e., inhibition of PIP2 hydrolysis by Opto-dHTH, localized Opto-dHTH induced RAW264.7 cell migration, dHTH does not interfere with G protein heterotrimer activation. Additionally, Opto-dHTH-induced adhered cell migration and suspended cell migration were conducted by three different experimenters. All experiments were repeated multiple times to test the reproducibility of the results. Results are analyzed from multiple cells and represented as mean±SEM. The exact number of cells used in the analysis is given in respective figure legends. Digital image analysis was performed using Andor iQ 3.1 software, and fluorescence intensity obtained from regions of interest was normalized to initial values (baseline). Data plot generation and statistical analysis were done using OriginPro software (OriginLab®). One-way ANOVA statistical tests were performed using OriginPro to determine the statistical significance between two or more populations of signaling responses. Tukey’s mean comparison test was performed at the p < 0.05 significance level for the one-way ANOVA statistical test. After obtaining the normalized data, PIP2 recovery rates were calculated using the NonLinear Curve Fitting tool in OriginPro. In the NonLinear Curve Fitting tool, each plot was fitted to DoseResp (dose-response) function under the pharmacology category by selecting the relevant range of data to be fitted. The mean values of hill slopes (P) obtained for each curve fitting are presented as the mean rates of PIP2 recovery.

### 3.7 Cytosolic calcium measurements

For intracellular calcium measurements, cells were cultured on cell culture dishes at 37 °C with 5% CO2. Experiments were performed 12–24 h after plating. Cells were washed twice with HBSS with calcium, pH 7.2, and incubated for 30 min at room temperature with the fluorescent calcium indicator Fluo-4 AM (2.28 μM). The fluorescence intensity of Fluo-4 AM was continuously imaged at 1s intervals using 515nm excitation and 542 nm emission with confocal microscopy. 488 nm excitation was not used to prevent the iLID activation on unintended occasions. Fluo-4 AM fluorescence intensities obtained from regions of interest were normalized to initial values.

### 3.8 Cell migration in isodense 3D Ficoll400 layer

RAW 264.5 cells were electroporated with GRPR, Opto-dHTH, and iLID CaaX using LONZA nucleofector following the manufacturer’s protocol. The cells were counted and resuspended in RAW media, plated on multiple 35 mm cell culture dishes (0.5 million cells per dish), and incubated at 37°C in 5% CO2 for 4 hours before imaging. Before imaging, cells were incubated with Versene EDTA for 2 min to detach them from the dish and spun for 3 minutes at 2800 rpm. Then cell pellet was resuspended in 40µL of 23% Ficoll400 (F8016; Sigma Aldrich) prepared in RAW cell media containing 1 µL of 0.1 mM bombesin. A 25 µL aliquot was pipetted from the cell suspension into a 4×5 mm glass cylinder (7030304; Bioptechs) fixed on a glass bottom dish with high-temperature silicon grease. A volume of 20 μl 10% Ficoll400 and 20 μl 5% Ficoll400 were added to create the desired gradient with the isodense layer with cells. Imaging was done 4-6 hours after electroporation.

## Supporting information

Ubeysinghe et al-Supporting Information

Subcellular PIP2 hydrolysis inhibition by OptodHTH

Subcellular OptodHTH recruitment

OptodHTH induced RAW 264.7 cell migration

## Abbreviations

GRPR: Gastrin releasing peptide receptor
PLC: Phospholipase C
PIP2: phosphatidylinositol 4,5-bisphosphate
IP3: inositol-1,4,5-triphosphate
DAG: diacylglycerol
GPCRs: G protein coupled receptors
GTP: Guanosine-5’-triphosphate
PKC: protein kinase C
Nt: N terminus
Ct: C terminus
PH: Pleckstrin homology
CTD: C-terminal domain
GEF: Guanine nucleotide exchange factor
GRK: G protein-coupled receptor kinase
ER: endoplasmic reticulum
MLCK: myosin light chain kinase
MLC: myosin light chain
MLCP: myosin light chain phosphatase
ROCK: Rho kinase

## Acknowledgment

We acknowledge Dr. N. Gautam (Washington University-School of Medicine, St. Louis, MO, USA) for providing us with plasmid DNA and RNAseq data. We also thank Dr. Zhou-Feng Chen (Washington University, St Louis, MO), Dr. Moritz Bunemann (Philipps-University Marburg, Germany), and Dr. Nevin Lambert (Augusta University, GA) for providing recombinant DNA plasmids. We thank the National Eye Institute for providing 11-cis-retinal. We also thank Ajith lab members Mithila Tennakoon, Senuri Piyawardena, Chathuri Rajarathne, and Adithya Chandu for various experimental support and discussions. We thank the Saint Louis University Institute for Drug and Biotherapeutic Innovation for providing computational resources and access to Schrödinger software with funding from the Saint Louis University Research Institute. We thank the Department of Biology at Saint Louis University for various instruments and infrastructure support.

## Author contribution

S.U. conducted the majority of experiments and performed the data analysis. D.K. generated the CRY2-mCherry-HTH construct and conducted the experiments. W.T. performed the RAW264.7 cell migrations experiments in Ficoll400 medium and modeled the structures using Alphafold2. D.W. performed the RAW264.7 cell migration experiment with melanopsin in the presence of Lyn-mRFP-dHTH and Lyn-mRFP. T.M. assisted with 3D cell migration experiments in Ficoll400. S.U. performed the statistical analysis. A.K., and S.U. conceptualized the project and wrote the manuscript.

## Data availability statement

The datasets used and analyzed during the current study are available from the corresponding author upon reasonable request.

