## Supplementary material for "Spatiotemporal optical control of Gαq-PLCβ interactions": Ubeysinghe et al-Supporting Information

#### Figure S1

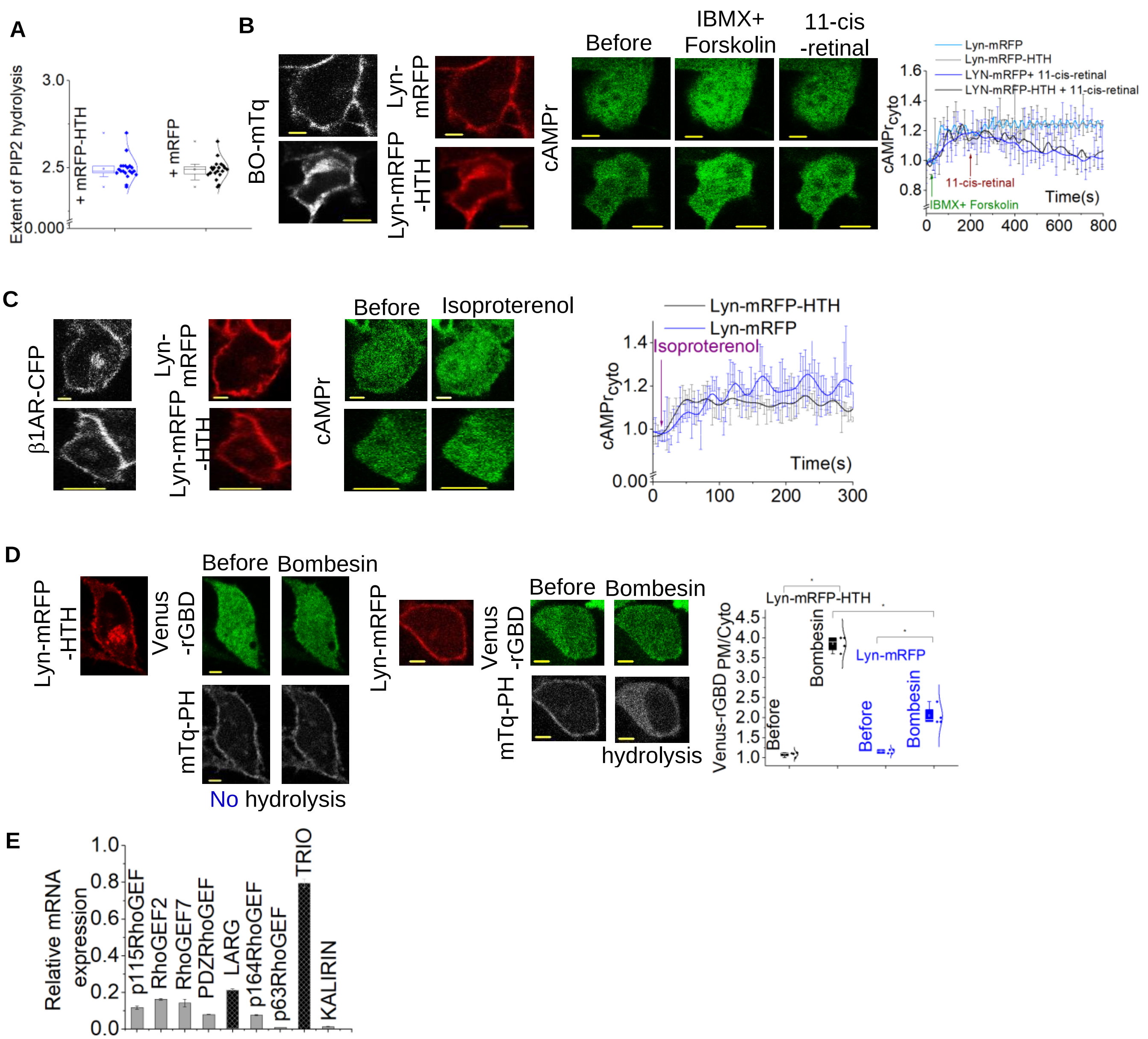

**Figure. S1:** **(A)** HeLa cells expressing cytosolic mRFP-HTH and the control cells with mRFP expression showed similar PIP2 hydrolysis responses. The whisker plot shows the extents of PIP2 hydrolysis under two conditions ($n \geq 18$). **(B)** HeLa cells expressing BO-mTq, cAMPr, Lyn-mRFP-HTH or Lyn-mRFP showed a similar initial cAMPr fluorescence increase upon addition of 10 μM forskolin in IBMX. This intensity was decreased upon addtion of 10μM 11-cis-retinal. However, in the control cells expressing the same DNA combinations that are not exposed to 11-cis-retinal, cAMPR fluorescence intensity remained at the higher level after the forskolin addition. The corresponding plot shows the cAMPr fluorescence in the cytosol (n = 6 for each condition). **(C)** HeLa cells expressing β1AR-CFP, cAMPr, and Lyn-mRFP-HTH or Lyn-mRFP exhibited a similar cAMPr fluorescence increase upon activation with 10 μM isoproterenol. The corresponding plot shows the cAMPr fluorescence in the cytosol (n= 10). **(D)** HeLa cells expressing GRPR, Lyn-mRFP-HTH, mTq-PH and Venus-rGBD showed a detectable Venus-rGBD recruitment to the plasma membrane upon the addition of 1 μM bombesin. However, these cells didn't show any PIP2 hydrolysis. The control cells expressing Lyn-mRFP showed a robust PIP2 hydrolysis and a minor Venus-rGBD recruitment to the plasma membrane, upon GRPR activation. The whisker plot shows the Venus-rGBD dynamics at the plasma membrane over time under the two conditions. **(E)** Relative mRNA expressions of different RhOGEFs in HeLa cells. HeLa cells show significant expression of LARG and TRIO compared to p63RhoGEF. Values are normalized to α4A-tubulin. Average curves were plotted using cells from ≥3 independent experiments. BO: blue opsin; β1AR: beta-1 adrenergic receptor; CFP: cyan fluorescence protein; mTq: mTurquoise; IBMX: 3-isobutyl-1-methylxanthine.

**Figure S2**

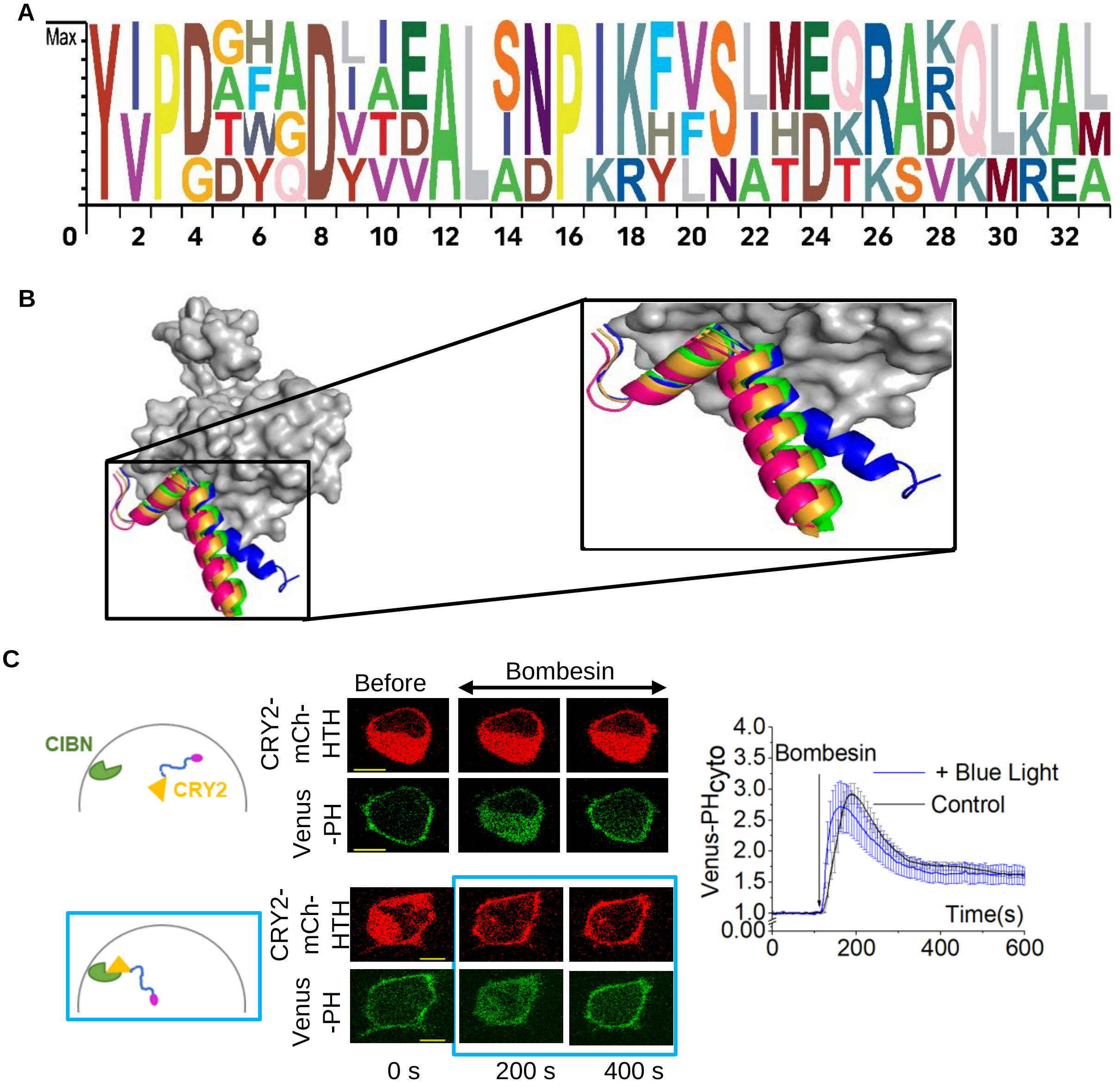


**Figure. S2: (A)** Position weight matrix showing the amino acid sequences alignment of the HTH domains of PLCβ 1-4 isoforms. **(B)** Modeled structure alignment of PLCβ1-4 isoforms with PLCβ3 (PDB ID: 4GNK). PLCβ1 HTH- green, PLCβ2 HTH- orange, PLCβ3 HTH- blue, PLCβ4 HTH- magenta. **(C)** HeLa cells expressing GRPR, Venus-PH, CRY2-mCh-HTH, and CIBN-CAAX exhibited significant PIP2 hydrolysis upon activation with 1 μM bombesin with and without optical activation. The plot shows the cytosolic PIP2 sensor dynamics with and without the optical command (n =15 with blue light and n= 20 without blue light). Average curves were plotted using cells from ≥3 independent experiments. The error bars represent SEM (standard error of mean). The scale bar = 5 μm. GRPR: Gastrin Releasing Peptide receptor; mCh: mCherry; PM: Plasma membrane; CRY2: Cryptochrome 2; PIP2: Phosphatidylinositol 4,5-bisphosphate; Cyto: cytosolic fluorescence; HTH: Helix-Turn-Helix; PH: Pleckstrin Homology. The blue box represents the blue light exposure.

### Figure S3

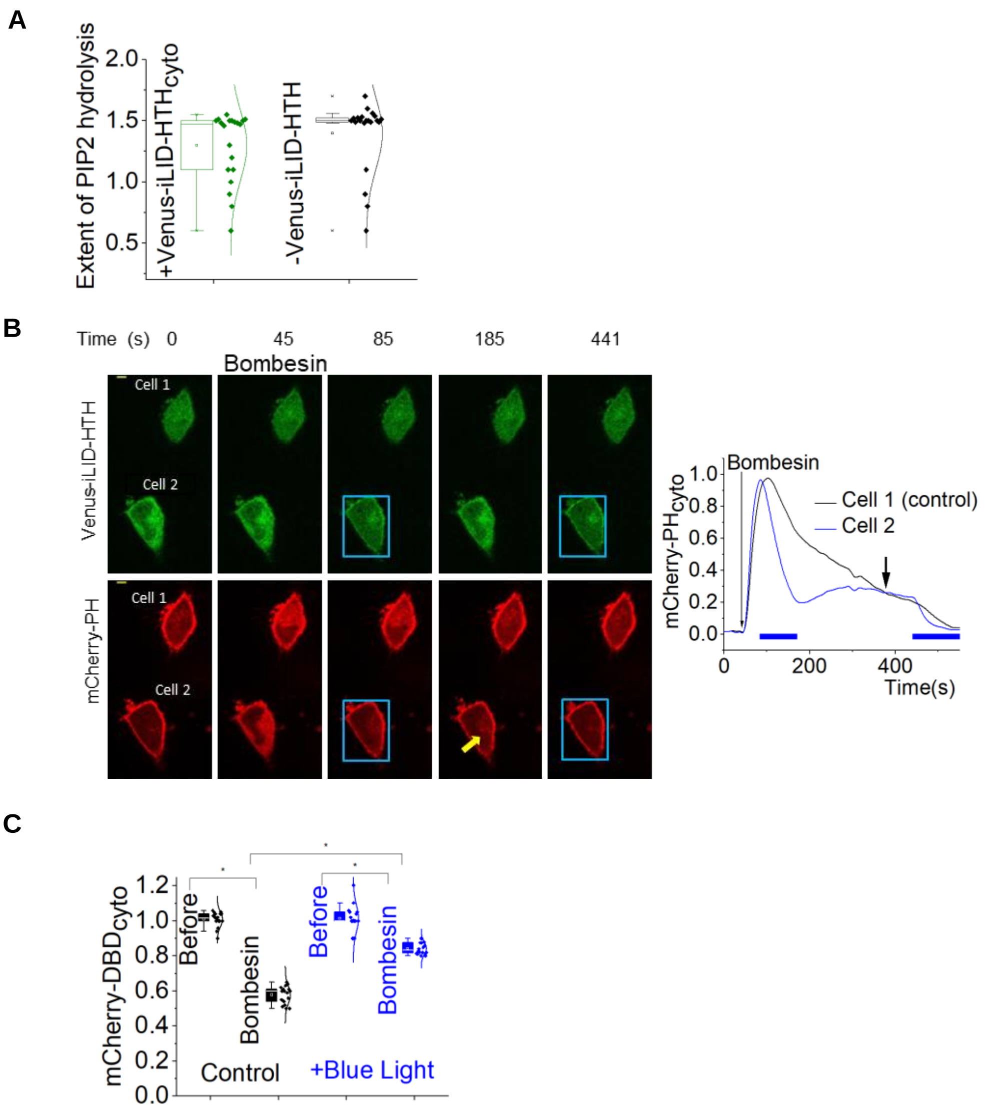


**Figure. S3: (A)** The whisker plot shows the extents of PIP2 hydrolysis in HeLa cells expressing cytosolic Venus-iLID-HTH, and control cells without Venus-ILID-HTH expression (n= 20 for each condition). **(B)** GRPR, Venus-iLID-HTH, Lyn-SspB and mCherry-PH were expressed in HeLa cells. GRPR was activated in two cells in the same field of vision. Venus-iLID-HTH was recruited to the plasma membrane in one cell while keeping the other cell intact. The PIP2 hydrolysis was inhibited only in blue light exposed cell. Blue lines on the plot: Blue light exposure on cell 2. Yellow arrow: The removal of the blue light induced PIP2 re-hydrolysis (n =2). The plot shows the PIP2 sensor dynamics in the cytosol. **(C)** The whisker box plot shows the mCherry-DBD Fluorescence intensity in the cytosol upon GRPR activation, with and without the optical command, compared to the basal level sensor fluorescence (n > 15 for each condition). Average curves were plotted using cells from ≥3 independent experiments. The error bars represent SEM (standard error of mean). The scale bar = 5 µm. The blue box represents the blue light exposure.

### Figure S4

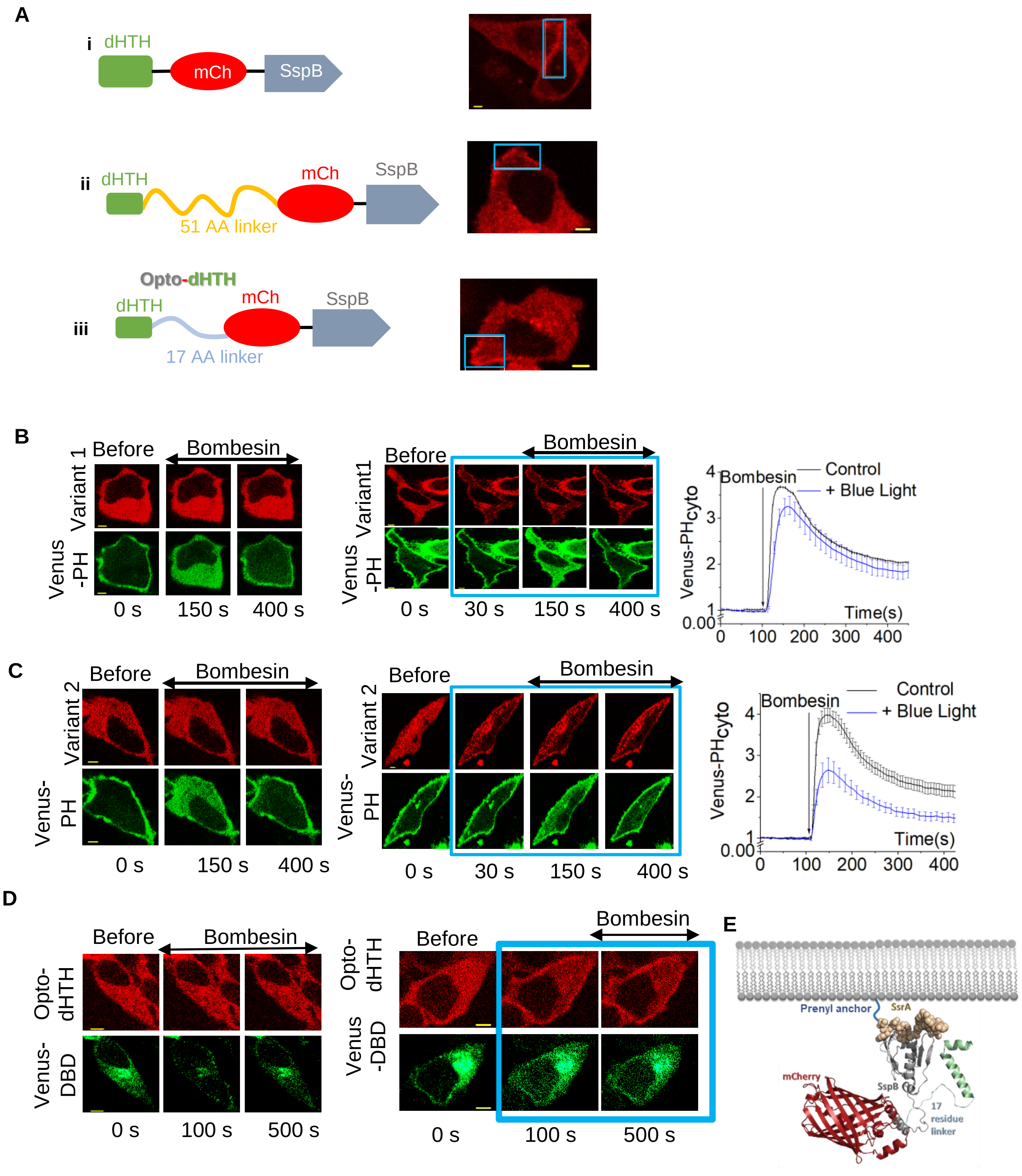
A
i
dHTH
mCh
SspB
ii
dHTH
51 AA linker
mCh
SspB
iii
Opto-dHTH
dHTH
17 AA linker
mCh
SspB
B
Before
Bombesin
Variant 1
Venus -PH
0 s
150 s
400 s
Before
Bombesin
Variant1
Venus -PH
0 s
30 s
150 s
400 s
Bombesin
Control
+ Blue Light
Venus-PHcyto
Time(s)
C
Before
Bombesin
Variant 2
Venus-PH
0 s
150 s
400 s
Before
Bombesin
Variant 2
Venus-PH
0 s
30 s
150 s
400 s
Bombesin
Control
+ Blue Light
Venus-PHcyto
Time(s)
D
Before
Bombesin
Opto-dHTH
Venus-DBD
0 s
100 s
500 s
Before
Bombesin
Opto-dHTH
Venus -DBD
0 s
100 s
500 s
E
Prenyl anchor
SsrA
mCherry
SspB
17 residue linker

**Figure. S4: (A) The modular design of the subcellular optogenetic GαqGTP inhibitor. (Ai)** First design with dHTH domain directly linked to mCherry which could be localized to a confined membrane region. **(Aii)** 51 residue linker was added between the dHTH domain and mCherry. This also could be recruited to a small membrane region. **(Aiii)** A truncated version of the second optogenetic module with 17 residues between the dHTH domain and mCherry also could be recruited to a confined membrane region. **(B)** Upon addition of 1 μM bombesin, HeLa cells expressing GRPR, Venus-PH, iLID-CAAX, and dHTH-mCh-SspB (variant 1) exhibited significant PIP2 hydrolysis with and without the optical command indicating its lack of inhibitory effect. The plot shows the cytosolic PIP2 sensor dynamics with and without the optical command (n =25 for each condition). **(C)** Activation of GRPR in HeLa cells expressing dHTH-51 residue linker-mCh-SspB (variant 2), iLID-CAAX, and Venus-PH resulted in partial inhibition of PIP2 hydrolysis under the optical command. The corresponding plot shows the cytosolic PIP2 sensor dynamics with and without the optical command (n = 30 for each condition). **(D)** Upon GRPR activation in HeLa cells expressing GRPR, Opto-dHTH, iLID-CAAX, and mCherry-DBD, induced a robust mCherry-DBD translocation to the plasma membrane, under no optical command. The mCherry-DBD translocation was not observed while Opto-dHTH is recruited to the plasma membrane using blue light (n =25 for each condition). **(E)** Modeled structure of Opto-dHTH using Alphafold2. Upon blue light exposure, Opto-dHTH interacts with membrane targeted iLID. Average curves were plotted using cells from ≥3 independent experiments. The error bars represent SEM (standard error of mean). The scale bar = 5 μm. The blue box represents the blue light exposure.

#### Figure S5

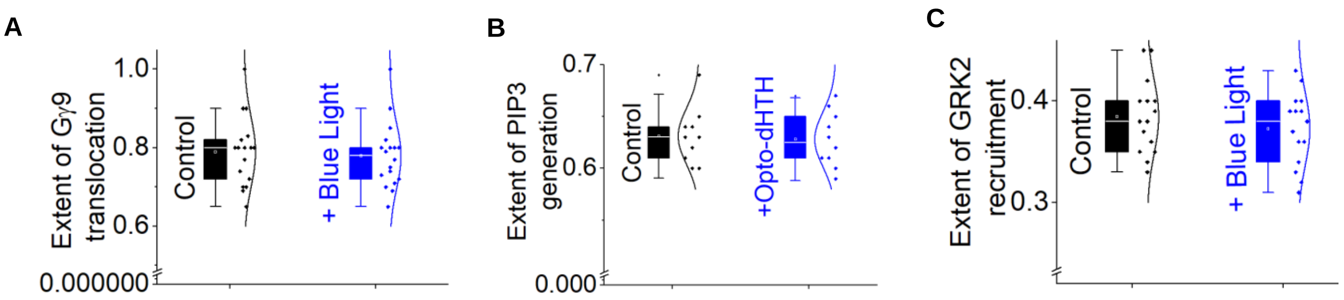


**Figure. S5: (A)** The whisker box plot shows the extent of Gγ9 translocation upon GRPR activation in the presence and absence of membrane recruited HTH (n ≥17 for each condition). **(B)** The whisker box plot shows the extent of PIP3 generation upon GRPR activation in RAW 264.7 cells in the presence and absence of membrane recruited HTH (n = 10 for each condition). **(C)** The whisker box plot shows the extent of GRK2- YFP membrane recruitment upon GRPR activation in HeLa cells in the presence and absence of membrane recruited HTH (n = 15 for each condition).

### Figure S6

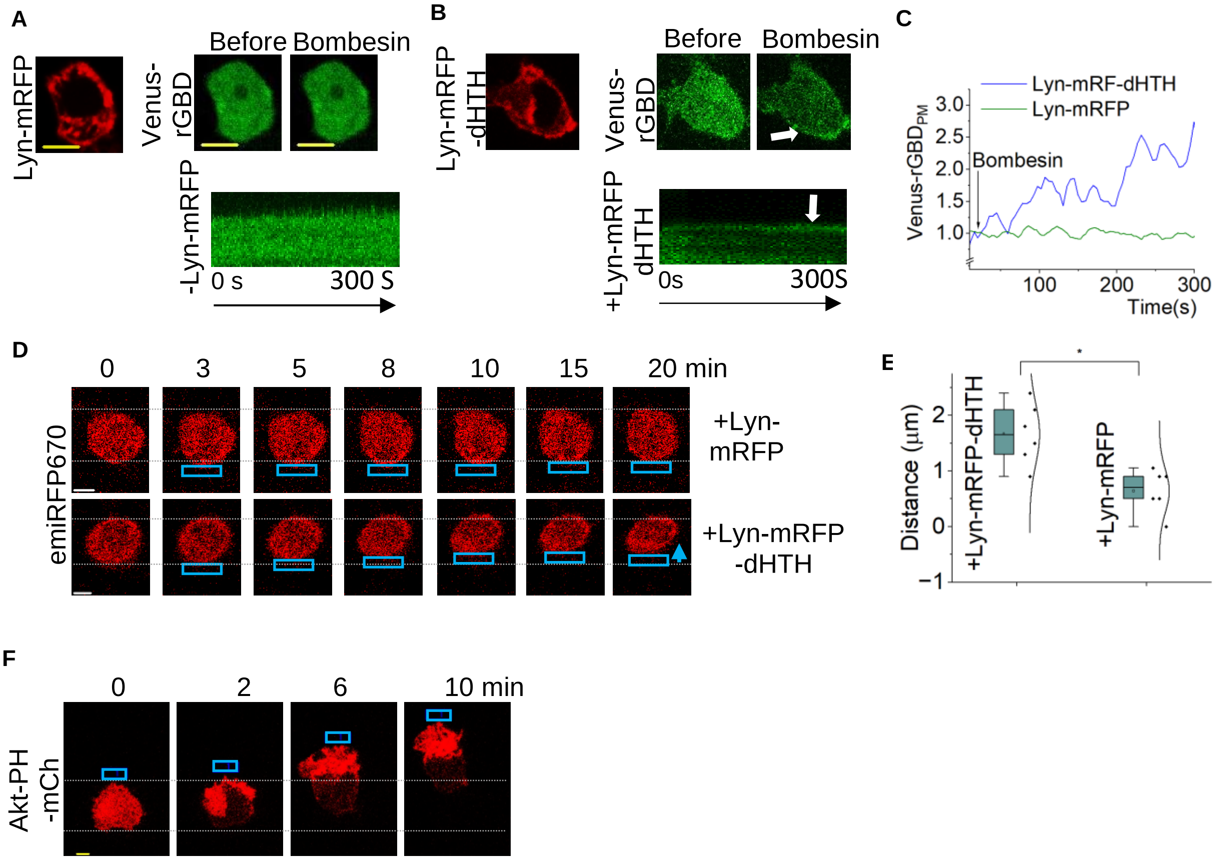


**Figure. S6: (A)** GRPR, Lyn-mRFP, and Venus-rGBD were expressed in RAW264.7 cells. Venus-rGBD didn't show a detectable recruitment to the plasma membrane of the cell. The kymograph shows the Venus-rGBD dynamics at the plasma membrane over time. The scale bar = 5 µm. **(B)** Similar experiment was performed in the presence of Lyn-mRFP-dHTH. Top: Upon GRPR activation, Venus-rGBD recruitment to the plasma membrane was observed (white arrow), indicating RhoA activation. Bottom: Kymographic view of the top cell showing Venus-rGBD accumulation at the plasma membrane over time. **(C)** The plot shows the dynamics of Venus-rGBD recruitment to the plasma membrane over time, with and without the expression of Lyn-mRFP-dHTH. **(D)** Top- The control RAW264.7 cells expressing melanopsin, emiRFP670 and Lyn-mRFP showed a barely detectable migration towards the opposite of localized melanopsin activation (Blue box indicates the localized melanopsin activation). In a similar experiment with Lyn-mRFP-dHTH, the cells showed significantly higher migration. **(E)** The whisker box plot shows the migration distances of the RAW264.7 cells within 10 min, upon localized Gαq coupled melanopsin activation in the presence of Lyn-mRFP or Lyn-mRFP-dHTH ($n \geq 5$ for each condition). **(F)** RAW264.7 cells expressing blue opsin-mTq and Akt-PH-mCherry showed a robust lamellipodia formation, PIP3 generation and migration towards the blue light exposure. The scale bar = 5 µm.

Table S1: Rationale design of Venus-iLID-HTH variants.

| Variant | Rationale | |
|---|---|---|
| HTH 36-mer | Complete HTH.<br>**YIPDDHQDYAEALINPIKHVSLMDQRARQLAALIGE** | 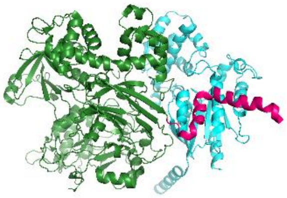 |
| HTH 31-mer | 5 residues from the Nt is deleted,assuming this will reduce the flexibility of the peptide, Strengthening the Gαq-HTH interactions.<br>**HQDYAEALINPIKHVSLMDQRARQLAALIGE** | 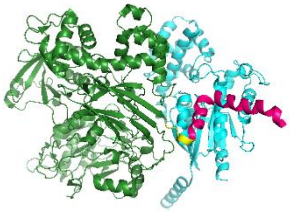 |
| HTH 27-mer | 5 residues from the Nt and 4 residues from the Ct are deleted, assuming the truncation increases the stability of the interactions between Gαq and HTH.<br>**HQDYAEALINPIKHVSLMDQRARQLAA** | 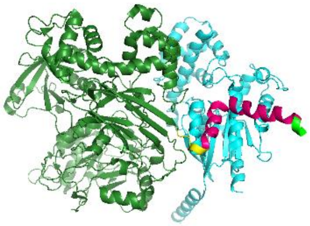 |
| HTH 27-mer I860A | Nt and Ct truncated HTH with I860A mutation.This mutation was previously shown to increase the phospholipase activity of PLCb3, and we assumed this substitution increases the affinity for Gαq.<br>**HQDYAEALANPIKHVSLMDQRARQLAA** | 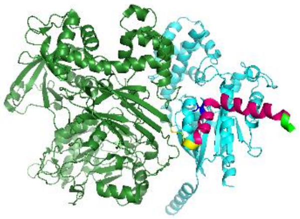 |
| HTH 27-mer P862H | Nt and Ct truncated HTH with P862H mutation. *Pro* involves forming the turn in protein secondary structures. *Pro* 862 was substituted with *His*, assuming this will disrupt the turn while changing its affinity for Gαq.<br>**HQDYAEALINHIKHVSLMDQRARQLAA** | 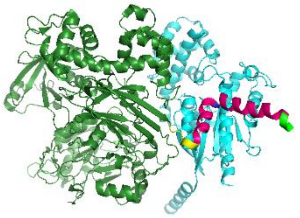 |
| HTH 22-mer | Ct of HTH is truncated to Shorten the peptide assuming this will form more stable interactions with Gαq.<br>**YIPDDHQDYAEALINPIKHVSL** | 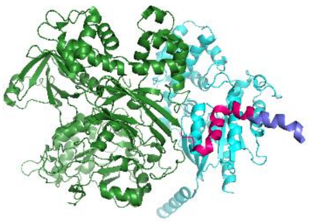 |

Table S2-A: One-way ANOVA statistics for extent of PIP2 hydrolysis with mRFP-HTH and mRFP.

| Descriptive Statistics | | | | |
|---|---|---|---|---|
| | N Analysis | Mean | Standard Deviation | SE of Mean |
| With mRFP-HTH | 20 | 2.493 | 0.0649 | 0.01451 |
| With mRFP | 20 | 2.4895 | 0.05591 | 0.0125 |

Table S2-B:

| Overall ANOVA | | | | | |
|---|---|---|---|---|---|
| | DF | Sum of Squares | Mean Square | F value | Prob>F |
| Model | 1 | 1.225E-4 | 1.225E-4 | 0.03339 | 0.85598 |
| Error | 38 | 0.13942 | 0.00367 | | |
| Total | 39 | 0.13954 | | | |

At the 0.05 level, the population means are not significantly different.

Table S3-A: One-way ANOVA statistics for the Venus-rGBD $_{PM/cyto}$ fluorescence before and after the GRPR activation in the presence of Lyn-mRFP-HTH or Lyn-mRFP.

| Descriptive Statistics | | | | |
|---|---|---|---|---|
| | N Analysis | Mean | Standard Deviation | SE of Mean |
| Before activation-Lyn-mRFP-HTH | 4 | 1.0825 | 0.0556 | 0.0278 |
| After activation-Lyn-mRFP_HTH | 4 | 3.85 | 0.19149 | 0.09574 |
| Before activation-Lyn-mRFP | 4 | 1.155 | 0.05196 | 0.02598 |
| After activation Lyn-mRFP | 4 | 1.675 | 0.17078 | 0.08539 |

Table S3-B:

| Overall ANOVA | | | | | |
|---|---|---|---|---|---|
| | DF | Sum of Squares | Mean Square | F value | Prob>F |
| Model | 3 | 20.27942 | 6.75981 | 377.5109 | <0.0001 |
| Error | 12 | 0.21487 | 0.01791 | | |
| Total | 15 | 20.49429 | | | |

At the 0.05 level, the population means are significantly different.

Table S4: Normalized RhoGEF expression profile in HeLa cells using RNA seq data.

| RhoGEF type | Expression (Number of reads) | a4A-tubulin expression (Number of reads) | Relative Expression (Normalized to α4Atubulin expression) |
|---|---|---|---|
| p115RhoGEF | 3311 | 19864 | 0.11712 |
| RhoGEF2 | 3158 | 19864 | 0.16315 |
| RhoGEF7 | 2580 | 19864 | 0.14275 |
| pDZRhoGEF | 1581 | 19864 | 0.07998 |
| LARG | 4847 | 19864 | 0.21234 |
| p164RhoGEF | 1451 | 19864 | 0.07657 |
| p63RhoGEF | 184 | 19864 | 0.009 |
| TRIO | 18411 | 19864 | 0.79630 |
| KALIRIN | 216 | 19864 | 0.01390 |

Table S5-A: One-way ANOVA statistics for the extent of PIP2 hydrolysis in cells with cytosolic Venus-iLID-HTH and without Venus-iLID-HTH expression.

| Descriptive Statistics | | | | |
|---|---|---|---|---|
| | N Analysis | Mean | Standard Deviation | SE of Mean |
| With cytosolic Venus-iLID-HTH | 20 | 1.2975 | 0.28525 | 0.06378 |
| Without Venus-iLID-HTH expression | 21 | 1.39681 | 0.28759 | 0.06276 |

Table S5-B:

| Overall ANOVA | | | | | |
|---|---|---|---|---|---|
| | DF | Sum of Squares | Mean Square | F value | Prob>F |
| Model | 1 | 0.10103 | 0.10103 | 1.23122 | 0.27396 |
| Error | 39 | 3.20019 | 0.08206 | | |
| Total | 40 | 3.30122 | | | |

At the 0.05 level, the population means are not significantly different.

Table S6-A: One-way ANOVA statistics for the cytosolic mCherry-DBD fluorescence before and after GRPR activation.

| Descriptive Statistics | | | | |
|---|---|---|---|---|
| | N Analysis | Mean | Standard Deviation | SE of Mean |
| Before activation-control | 18 | 1.009 | 0.04126 | 0.00973 |
| After activation-control | 18 | 0.57824 | 0.0499 | 0.0121 |
| Before activation-with blue light | 16 | 1.00825 | 0.04383 | 0.01096 |
| After activation with blue light | 16 | 0.84331 | 0.03427 | 0.00857 |

Table S6-B:

| Overall ANOVA | | | | | |
|---|---|---|---|---|---|
| | DF | Sum of Squares | Mean Square | F value | Prob>F |
| Model | 3 | 2.10538 | 0.70179 | 383.72078 | <0.0001 |
| Error | 63 | 0.11522 | 0.00183 | | |
| Total | 66 | 2.2206 | | | |

At the 0.05 level, the population means are significantly different.

Table S7-A: One-way ANOVA statistics for the relative Fluo4-AM fluorescence increase before and after the GRPR activation.

| Descriptive Statistics | | | | |
|---|---|---|---|---|
| | N Analysis | Mean | Standard Deviation | SE of Mean |
| Before activation-control | 20 | 0.98925 | 0.03471 | 0.00776 |
| After activation-control | 20 | 2.943 | 0.49767 | 0.11128 |
| Before activation-with blue light | 20 | 1.06188 | 0.23199 | 0.05188 |
| After activation with blue light | 20 | 1.4735 | 0.18325 | 0.04098 |

Table S7-B:

| Overall ANOVA | | | | | |
|---|---|---|---|---|---|
| | DF | Sum of Squares | Mean Square | F value | Prob>F |
| Model | 3 | 49.62195 | 16.54065 | 196.74722 | <0.0001 |
| Error | 76 | 6.38936 | 0.08407 | | |
| Total | 79 | 56.01131 | | | |

At the 0.05 level, the population means are significantly different.

Table S8-A: One-way ANOVA statistics for extent of PIP2 hydrolysis with Venus-iLID-HTH and Venus-iLID-dHTH.

| Descriptive Statistics | | | | |
|---|---|---|---|---|
| | N Analysis | Mean | Standard Deviation | SE of Mean |
| With Venus-iLID-HTH | 20 | 0.2373 | 0.03079 | 0.00688 |
| With Venus-iLID-dHTH | 20 | 0.051 | 0.03463 | 0.00774 |

Table S8-B:

| Overall ANOVA | | | | | |
|---|---|---|---|---|---|
| | DF | Sum of Squares | Mean Square | F value | Prob>F |
| Model | 1 | 0.34708 | 0.34708 | 323.27216 | 0 |
| Error | 38 | 0.0408 | 0.00107 | | |
| Total | 39 | 0.38788 | | | |

At the 0.05 level, the population means are significantly different.

Table S9-A: One-way ANOVA statistics for the relative Fluo4-AM fluorescence increase before and after the GRPR activation.

| Descriptive Statistics | | | | |
|---|---|---|---|---|
| | N Analysis | Mean | Standard Deviation | SE of Mean |
| Before activation-control | 25 | 1.0524 | 0.1712 | 0.03424 |
| After activation-control | 25 | 2.81483 | 0.39202 | 0.0784 |
| Before activation-with blue light | 25 | 1.0236 | 0.12359 | 0.02472 |
| After activation with blue light | 25 | 1.19842 | 0.06675 | 0.01335 |

Table S9-B:

| Overall ANOVA | | | | | |
|---|---|---|---|---|---|
| | DF | Sum of Squares | Mean Square | F value | Prob>F |
| Model | 3 | 56.12606 | 18.70869 | 369.15978 | <0.0001 |
| Error | 96 | 4.86519 | 0.05068 | | |
| Total | 99 | 60.99126 | | | |

At the 0.05 level, the population means are significantly different.

Table S10-A: One-way ANOVA statistics for Gγ9 translocation in HeLa cells.

| Descriptive Statistics | | | | |
|---|---|---|---|---|
| | N Analysis | Mean | Standard Deviation | SE of Mean |
| Control | 17 | 0.78941 | 0.08835 | 0.02143 |
| With Blue light | 18 | 0.77889 | 0.0826 | 0.01947 |

Table S10-B:

| Overall ANOVA | | | | | |
|---|---|---|---|---|---|
| | DF | Sum of Squares | Mean Square | F value | Prob>F |
| Model | 1 | 9.68105E-4 | 9.68105E-4 | 0.13263 | 0.71804 |
| Error | 33 | 0.24087 | 0.0073 | | |
| Total | 34 | 0.24184 | | | |

At the 0.05 level, the population means are not significantly different.

Table S11-A: One-way ANOVA statistics for PIP3 generation in RAW 264.7 cells.

| Descriptive Statistics | | | | |
|---|---|---|---|---|
| | N Analysis | Mean | Standard Deviation | SE of Mean |
| Without Opto-dHTH | 10 | 0.631 | 0.02961 | 0.00936 |
| With Opto-dHTH | 10 | 0.64 | 0.03232 | 0.01022 |

Table S11-B:

| Overall ANOVA | | | | | |
|---|---|---|---|---|---|
| | DF | Sum of Squares | Mean Square | F value | Prob>F |
| Model | 1 | 4.05E-4 | 4.05E-4 | 0.42163 | 0.52432 |
| Error | 18 | 0.01729 | 9.60556E-4 | | |
| Total | 19 | 0.0177 | | | |

At the 0.05 level, the population means are not significantly different.

Table S12-A: One-way ANOVA statistics for GRK2-YFP recruitment in HeLa cells.

| Descriptive Statistics | | | | |
|---|---|---|---|---|
| | N Analysis | Mean | Standard Deviation | SE of Mean |
| Control | 15 | 0.38467 | 0.03662 | 0.00945 |
| With Blue light | 15 | 0.37267 | 0.03575 | 0.00923 |

Table S12-B:

| Overall ANOVA | | | | | |
|---|---|---|---|---|---|
| | DF | Sum of Squares | Mean Square | F value | Prob>F |
| Model | 1 | 0.00108 | 0.00108 | 0.82473 | 0.37155 |
| Error | 28 | 0.03667 | 0.00131 | | |
| Total | 29 | 0.03775 | | | |

At the 0.05 level, the population means are not significantly different.

Table S13-A: One-way ANOVA statistics for suspended and adhered RAW264.7 cell migrations, upon GRPR activation.

| Descriptive Statistics | | | | |
|---|---|---|---|---|
| | N Analysis | Mean | Standard Deviation | SE of Mean |
| Suspended cells | 10 | 4.75 | 0.29533 | 0.09339 |
| Adherent cells | 5 | 12.14 | 0.29665 | 0.13266 |

Table S13-B:

| Overall ANOVA | | | | | |
|---|---|---|---|---|---|
| | DF | Sum of Squares | Mean Square | F value | Prob>F |
| Model | 1 | 182.04033 | 182.04033 | 2081.37584 | <0.0001 |
| Error | 13 | 1.137 | 0.08746 | | |
| Total | 14 | 183.17733 | | | |

At the 0.05 level, the population means are significantly different.

Table S14A: One-way ANOVA statistics for Trailing Edge retraction of RAW264.7 cells

| Descriptive Statistics – Trailing Edge | | | | |
|---|---|---|---|---|
| | N Analysis | Mean | Standard Deviation | SE of Mean |
| Control | 9 | 4.75556 | 0.31667 | 0.10556 |
| Gallein | 10 | 4.59 | 0.1792 | 0.05667 |
| Wortmannin | 9 | 4.74444 | 0.24037 | 0.08012 |
| YM-254890 | 9 | 0.14111 | 0.04512 | 0.01504 |
| Y-27632 | 9 | 0.14444 | 0.04362 | 0.01454 |

Table S14-B:

| Overall ANOVA- Trailing edge | | | | | |
|---|---|---|---|---|---|
| | DF | Sum of Squares | Mean Square | F value | Prob>F |
| Model | 4 | 223.72352 | 55.93088 | 1446.833 | <0.0001 |
| Error | 41 | 1.58496 | 0.03866 | | |
| Total | 45 | 225.30848 | | | |

Table S15A: One-way ANOVA statistics for RAW264.7 cell migrations upon melanopsin activation in the presence of Lyn-mRFP or Lyn-mRFP-dHTH.

| Descriptive Statistics | | | | |
|---|---|---|---|---|
| | N Analysis | Mean | Standard Deviation | SE of Mean |
| With Lyn-mRFP | 6 | 1.66667 | 0.5465 | 0.22311 |
| With Lyn-mRFP-dHTH | 6 | 0.64167 | 0.38784 | 0.15833 |

Table S15-B:

| Overall ANOVA | | | | | |
|---|---|---|---|---|---|
| | DF | Sum of Squares | Mean Square | F value | Prob>F |
| Model | 1 | 3.15188 | 3.15188 | 14.03693 | 0.00381 |
| Error | 10 | 2.24542 | 0.22454 | | |
| Total | 11 | 5.39729 | | | |

At the 0.05 level, the population means are significantly different.

#### SUPPLEMENTARY MOVIE LEGENDS

**Movie S1.** Localized optical recruitment of Opto-dHTH using 445 nm light pulses (white box).

**Movie S2.** Subcellular PIP2 hydrolysis inhibition resulted by the localized Opto-dHTH recruitment. White box indicates the blue light exposed region of the cell.

**Movie S3.** Localized optical of recruitment of Opto-dHTH in RAW264.7 cells results in cell migration opposite to the blue light exposure. White box indicates the blue light exposed region of the cell.
